# Secondary structure transitions and dual PIP2 binding define cardiac KCNQ1-KCNE1 channel gating

**DOI:** 10.1101/2025.06.10.657059

**Authors:** Ling Zhong, Xiaoqing Lin, Xinyu Cheng, Shuangyan Wan, Yaoguang Hua, Weiwei Nan, Bin Hu, Zhenzhen Yan, Dexiang Jiang, Hangyu Zhang, Fengjiao Liu, Chenxin Xiao, Zhuo Zhou, Haijie Yu, Lijuan Ma, Chen Huang, Vincent Kam Wai Wong, Sookja Kim Chung, Bing Shen, Zhi-Hong Jiang, Erwin Neher, Wandi Zhu, Jin Zhang, Panpan Hou

**Affiliations:** School of Pharmacy, Faculty of Medicine & State Key Laboratory of Quality Research in Chinese Medicines, Dr. Neher’s Biophysics Laboratory for Innovative Drug Discovery, Macau University of Science and Technology, Macau SAR, China. Macau University of Science and Technology Innovation Technology Research Institute. Hengqin, Guangdong, China; The MOE Basic Research and Innovation Center for the Targeted Therapeutics of Solid Tumors, The Second Affiliated Hospital, Jiangxi Medical College, Nanchang University, Nanchang, China; School of Basic Medical Sciences, Jiangxi Medical College, Nanchang University, Nanchang, China; School of Basic Medical Sciences, Gannan Medical University, Ganzhou, China; College of Medicine, Department of Molecular Medicine and Therapeutics, The Ohio State University. Columbus, OH, United States

## Abstract

The KCNQ1+KCNE1 potassium channel complex forms the slow delayed rectifier current (I_Ks_) critical for cardiac repolarization. Loss-of-function variants in *KCNQ1* and *KCNE1* cause long QT syndrome types 1 and 5 (LQT1/LQT5), accounting for over one-third of clinical LQTS cases. Despite prior structural work on KCNQ1 and KCNQ1+KCNE3, the structural basis of KCNQ1+KCNE1 remains unresolved. Using cryo-EM and electrophysiology, we determined high-resolution (2.5-3.4 Å) structures of human KCNQ1+KCNE1 in both closed and open states. KCNE1 occupies a pivotal position at the interface of three KCNQ1 subunits, inducing seven “helix-to-loop” transitions in KCNQ1 transmembrane segments. These structural rearrangements: 1) stabilize the closed pore and the conformation of the intermediate voltage-sensing domain, thereby determining channel gating, ion permeation, and single channel conductance; 2) enable a dual-PIP2 modulation mechanism, where one PIP2 occupies the canonical site, while the second PIP2 bridges the S4-S5 linker, KCNE1, and the adjacent S6’, stabilizing channel opening; 3) create a fenestration capable of binding compounds specific for KCNQ1+KCNE1 (e.g., AC-1). Together, these findings reveal a previously unrecognized large-scale secondary structural transition during ion channel gating that fine-tunes I_Ks_ function and provides a foundation for targeted LQTS therapy development.

## Introduction

The KCNQ1+KCNE1 channel complex serves as a master regulator of cardiac repolarization and electrical stability of the heart ^1-4^. Dysfunctional variants in either the pore-forming subunit KCNQ1 or the auxiliary subunit KCNE1 are directly implicated in long QT syndrome types 1 and 5 (LQT1 and LQT5), two leading causes of sudden cardiac death in young individuals ^5-7^. The cardiac-specific KCNE1 auxiliary subunit (also known as MinK) is a single transmembrane protein consisting of 129 amino acids ^1-4,8^. When co-assembled with KCNQ1 to form the KCNQ1+KCNE1 (I_Ks_) channel ^3,4,9^, this heteromeric complex dynamically orchestrates ventricular action potential termination through three key features: 1) Slow-activating gating kinetics. KCNE1 significantly increases macroscopic current amplitude and single channel conductance, slows activation and deactivation kinetics, and right-shifts the voltage-dependent activation ^10-17^; 2) Altered ion selectivity. KCNE1 reduces the rubidium/potassium (Rb^+^/K^+^) permeability ratio through the selectivity filter ^16,18-21^; and 3) Changed pharmacological profile. KCNE1 modulates the channel’s sensitivity to activators (e.g. ML277, polyunsaturated fatty acids) and blockers (e.g. XE991, AC-1) ^16,18,22-29^. Despite decades of functional electrophysiology characterization, the structural basis for KCNE1’s profound modulation of KCNQ1 remains elusive. Addressing this gap is critical for revealing the mechanisms underlying I_Ks_ channel allosteric gating and enabling structure-guided therapeutics development for LQTS.

The KCNQ1 channel (K_V_7.1/K_V_LQT1) is a voltage-gated potassium channel ^3,4,9^, with a canonical tetrameric architecture ^30-34^. Each subunit comprises six transmembrane segments (S1-S6), organized into four peripheral voltage-sensing domains (VSD, S1-S4) and a central pore domain (PD, S5-S6). The VSDs are connected to the PD via S4-S5 linkers. KCNQ1 exhibits a domain-swapped architecture where each VSD is in close contact with a neighboring subunit’s PD ^30-34^. Activation of KCNQ1 follows a "Hand-and-Elbow" gating mechanism involving sequential VSD to pore transitions ^23^: 1) S4 movement from resting to intermediate state triggers intra-subunit "hand" interactions (between S4-S5 linker and lower S6), producing an intermediate open (IO) state; 2) subsequent S4 transition to activated state engages inter-subunit "elbow" interactions (between the S4/S4-S5 linker and the neighboring pore), yielding the activated open (AO) state ^11,16,18,23,30,35-37^. Notably, the KCNE1 association selectively suppresses the IO state but enhances the AO state, restricting the KCNQ1+KCNE1 channel opens only to the AO state ^16,18,23,30,36,37^. Although this allosteric regulation is functionally defined, it lacks structural evidence, which leaves several key questions unresolved. How does a small KCNE1 subunit completely reshape KCNQ1’s conformational changes during gating? What structural determinants underlie I_Ks_’s distinct function from KCNQ1 alone and the epithelial KCNQ1+KCNE3 channel that is constitutively open ^8,10,13,35,38,39^?

In this study, we integrated cryo-electron microscopy (cryo-EM) and electrophysiology to elucidate the structural basis of KCNE1’s regulation of KCNQ1 that shapes cardiac repolarization. We determined high-resolution (2.5-3.4 Å) structures of the human KCNQ1 in apo state (closed), KCNQ1+KCNE1 complexes in apo state (closed), and PIP2-bound open state. Detailed analyses reveal that KCNE1 occupies a strategic position at the interface of three KCNQ1 subunits, inducing seven secondary structural “helix-to-loop” transitions in KCNQ1 transmembrane segments, which: 1) stabilize the closed pore and intermediate VSD state, thereby determining channel gating, ion permeation, and single channel conductance; 2) establish a dual-PIP2 modulation mechanism, where one PIP2 occupies the canonical site, while the second PIP2 bridges the S4-S5 linker, KCNE1, and adjacent S6’, stabilizing channel opening; 3) create a fenestration that allows binding of compounds to selectively modulate KCNQ1+KCNE1 activity. Taken together, our findings provide a new structural paradigm for ion channel gating, and establish a structural framework for developing targeted therapies against LQT syndromes.

## Results

### Structure of the KCNQ1+KCNE1 channel

The KCNQ1 activation currents can be fitted by a double exponential function, with the fast (*τ*_-f_) and slow (*τ*_-s_) components representing the currents of the IO and AO states, respectively ^11,18,23,30,35-37^ (**Figure 1A**). The association of KCNE1 profoundly modulates KCNQ1 channel gating properties, which: 1) increases the current amplitude, along with slowed activation kinetics (**Figure 1A**); 2) right-shifts the voltage-dependent activation by ∼70 mV (**Figure 1C**); 3) suppresses the IO current, so that the KCNQ1+KCNE1 channel opens only to the slow-activating AO state (**Figure 1A**).

**Figure 1.**
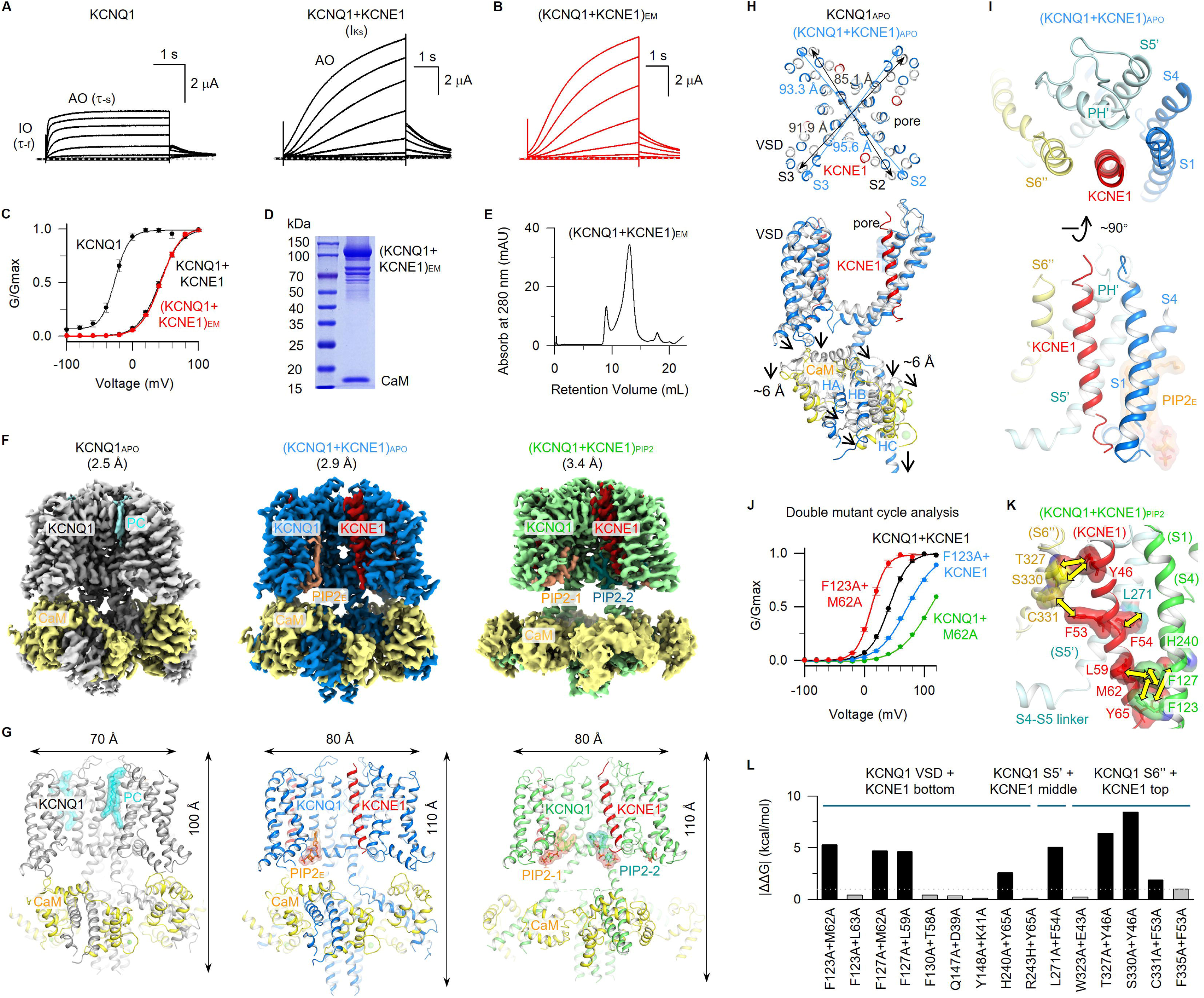
Overall closed- and open-state structures of KCNQ1+KCNE1, and direct interactions between KCNQ1 and KCNE1. **(A)** Representative activation currents of KCNQ1 and KCNQ1+KCNE1 recorded from -120 mV to +80 mV for 4 s or 5 s, and then returned to -40 mV for tail currents. KCNQ1 currents show fast and slow components (*τ*_-f_, *τ*_-s_), approximating currents of IO and AO states. **(B)** Representative activation currents of (KCNQ1+KCNE1)_EM_. **(C)** G–V relationships of KCNQ1, KCNQ1+KCNE1, and (KCNQ1+KCNE1)_EM_ (red). **(D)** SDS-PAGE analysis of the (KCNQ1+KCNE1)_EM_ sample. **(E)** Size-exclusion chromatography of (KCNQ1+KCNE1)_EM_ on Superose 6. **(F)** Side view of KCNQ1 density maps in the apo condition, and (KCNQ1+KCNE1)_EM_ density maps in apo and in the presence of PIP2. An endogenous phosphatidylcholine (PC) molecule was observed in KCNQ1_APO_, and an endogenous PIP2 molecule was observed in (KCNQ1+KCNE1)_APO_; while 2 PIP2 molecules, PIP2-1 and PIP2-2, were observed in each subunit of (KCNQ1+KCNE1)_PIP2_. Color code: KCNQ1_APO_ (gray), (KCNQ1+KCNE1)_APO_ (blue), (KCNQ1+KCNE1)_PIP2_ (green), KCNE1 (red), PIP2-1 (orange), PIP2-2 (dark green), and CaM (yellow). **(G)** Structure models of KCNQ1_APO_, (KCNQ1+KCNE1)_APO_, and (KCNQ1+KCNE1)_PIP2_. **(H)** KCNE1-induced horizontal expansion to the VSD (from 85.1 Å to 93.3 Å for S2-I161, and from 91.9 Å to 95.6 Å for S3-C214), and vertical expansion to the S2-S3 linker and the cytosolic domain (∼6 Å). Structures were aligned to the filter. **(I)** Top and side views of interfaces between KCNQ1 and KCNE1. **(J)** Double mutant cycle analysis to test the interaction between KCNQ1-F123 and KCNE1-M62 with ΔΔ*G* =5.3 kcal/mol. **(K-L)** Summary of KCNQ1 and KCNE1 interactions at different parts tested by double mutant cycle analysis. ΔΔ*G* >1 kcal/mol are: F123 and M62 (5.3 kcal/mol), F127 and M62 (4.7 kcal/mol), F127 and L59 (4.6 kcal/mol), H240 and Y65 (2.6 kcal/mol), L271 and F54 (5.0 kcal/mol), T327 and Y46 (8.4 kcal/mol), S330 and Y46 (6.4 kcal/mol), C331 and F53 (1.9 kcal/mol).

To elucidate the structural basis of these intensive KCNE1 modulations, we determined cryo-EM structures of human KCNQ1+KCNE1 channel under different conditions. Initial attempts with full-length human KCNQ1 co-expressed with KCNE1 yielded only KCNQ1 protein lacking KCNE1 after purification (**Figure S1**), likely due to the structural flexibility of the N- and C-termini of KCNQ1 ^31,32^. We therefore engineered a stabilized construct (KCNQ1+KCNE1)_EM_, fusing KCNE1 to N/C-termini truncated KCNQ1 (KCNE1-ΔN/ΔC/KCNQ1). A similar strategy has been widely used in both structural and functional studies. This (KCNQ1+KCNE1)_EM_ construct produced channels exhibiting almost identical current kinetics and G–V relation to the wild-type (WT) KCNQ1+KCNE1 channel (**Figure 1B,C**). SDS-page results showed that the (KCNQ1+KCNE1)_EM_ protein was successfully purified with size exclusion chromatography (**Figure 1D,E**).

We determined high resolution structures of human KCNQ1 in the apo state: KCNQ1_APO_ (2.5 Å), and KCNQ1+KCNE1 complexes in both the apo state: (KCNQ1+KCNE1)_APO_ (2.9 Å) and in the presence of PIP2: (KCNQ1+KCNE1)_PIP2_ (3.4 Å) (**Figure 1F,G, Figures S2-S4**). All structures were determined with C4 symmetry. The KCNQ1_APO_ structure, serving as a reference structure, aligned well with previous studies ^32^, but revealed a previously unrecognized lipid molecule bound in the cleft formed by three different subunits at the extracellular side (cyan, **Figure 1F,G, Figure S2**). The EM map resolved this lipid density to match phosphatidylcholine (PC) (**Figure 1F,G, Figure S2**), the predominant phospholipid in the outer leaflet of the plasma membrane ^40^. Interestingly, in both (KCNQ1+KCNE1)_APO_ and (KCNQ1+KCNE1)_PIP2_ structures, the KCNE1 transmembrane segment (D39-I66 for (KCNQ1+KCNE1)_APO_, and D39-S68 for (KCNQ1+KCNE1)_PIP2_) occupies the same cleft, in place of the PC molecule (**Figure 1F-I**). The remaining regions of KCNE1 were not resolved, presumably also due to their structural flexibility.

The KCNQ1+KCNE1 structure adopts a domain-swapped architecture (**Figure 1G**, **Figure S5**). Each tetrameric channel can bind four KCNE1 subunits, following a KCNQ1:KCNE1 stoichiometry of 4:4 (**Figure 1F,G**). The 4:4 stoichiometry was confirmed by symmetry-free refinement (**Figure S6**). Structural analysis of KCNQ1_APO_and (KCNQ1+KCNE1)_APO_, aligned to the selectivity filter, shows that the association of KCNE1 induced a horizontal expansion of the VSDs (e.g. S2-I161: 85.1→93.3 Å; S3-C214: 91.9→95.6 Å), and a downward movement to the S2-S3 linker, the S4-S5 linker, and the entire cytosolic domain by ∼6 Å (**Figure 1H**). Consequently, KCNE1 induced a global structural expansion to almost every amino acid of KCNQ1.

### Double mutant cycle analysis mapped interactions between KCNQ1 and KCNE1

The positioning of KCNE1 allows it to directly interact with the VSD (S1 and S4, bule) of one subunit, the pore domain (S5’ and the pore helix’, PH’, green) of an adjacent subunit, and the pore domain (S6’’, yellow) of the opposing subunit (**Figure 1I**). This tripartite binding geometry enables KCNE1 to simultaneously interact with multiple functional modules of KCNQ1.

To systematically map molecular interactions between KCNQ1 and KCNE1, we performed double mutant cycle (DMC) analysis ^12,41-43^. Taking the spatially proximal residue pair KCNQ1-F123 (S1) and KCNE1-M62 as an example (**Figure 1J-L**), we measured activation energies of WT KCNQ1+KCNE1 (*G_0_*= 3.6 kcal/mol), F123A+KCNE1 (*G_1_*= 4.3 kcal/mol), KCNQ1+M62A (*G_2_*= 5.9 kcal/mol), and F123A+M62A (*G*= 1.3 kcal/mol). The total energy change ΔΔ*G*= (*G_1_*+*G_2_*)-(*G_0_*+*G*)= 5.3 kcal/mol (>1 kcal/mol), confirming their direct interaction (**Figure 1J**).

Through systematic DMC analysis, we identified seven additional interaction pairs between KCNE1 and all three contacting KCNQ1 subunits (**Figure 1K,L**, **Figure S7**): 1) S1 (F127) with bottom of KCNE1 (M62: 4.7 kcal/mol; L59: 4.6 kcal/mol), and S4 (H240) directly interacts with KCNE1-Y65 (2.6 kcal/mol); 2) S5’ (L271) with middle of KCNE1-F54 (5.0 kcal/mol), S6’’ (C331) with KCNE1-F53 (1.9 kcal/mol); and 3) S6’’ (T327 and S330) with top of KCNE1-Y46 (8.4 and 6.4 kcal/mol, respectively). Control experiments identified seven non-interacting residue pairs despite their physical proximity (F123/L63, F130/T58, Q147/D39, Y148/K41, R243/Y65, W323/E43, and F335/F53 showed ΔΔ*G* <1 kcal/mol; **Figure 1K**, **Figure S8**), and another six mutant pairs with non-functional I_Ks_ currents that precluded DMC analysis (Y267/F54, Y267/T58, L303/Y46, W304/Y46, V307/Y46, W323/Y46, **Figure S9**). These results map KCNE1’s extensive interaction network with all three KCNQ1 subunits, which is necessary for its global structural and functional modulations of KCNQ1.

### KCNE1 stabilizes the intermediate VSD state of KCNQ1

The voltage-dependent activation of KCNQ1+KCNE1 channels proceeds through three sequential steps: VSD activation, VSD-pore coupling, and pore opening. We next aim to elucidate the structural basis of KCNQ1+KCNE1 gating process.

To document KCNE1-induced conformational changes to VSD activation, we aligned (KCNQ1+KCNE1)_APO_ and (KCNQ1+KCNE1)_PIP2_ structures to the selectivity filter of KCNQ1_APO_. Remarkably, the KCNE1 association induces a significant ∼12° rotation (counterclockwise from top view) of the VSD relative to the pore (**Figure 2A**), and the S1-S4 segment thus moves ∼10 Å (**Figure S10**). These dramatic changes may alter the VSD activation.

**Figure 2.**
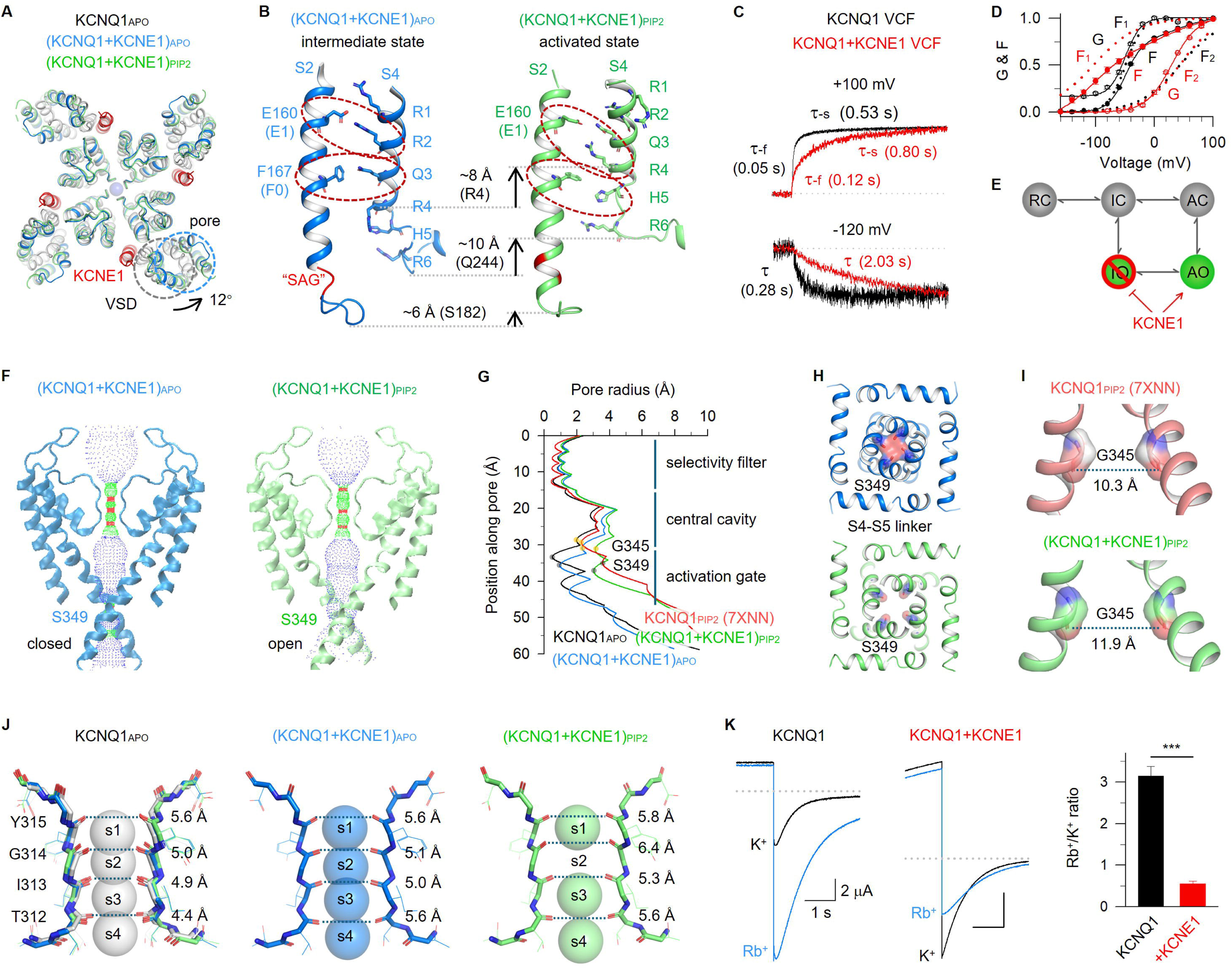
KCNE1-induced structural changes to the KCNQ1 VSD and pore. **(A)** Structural comparison of KCNQ1_APO_, (KCNQ1+KCNE1)_APO_, and (KCNQ1+KCNE1)_PIP2_ to show KCNE1 indued a ∼12° rotation to the VSD (counterclockwise). **(B)** Structural comparison of VSD states between (KCNQ1+KCNE1)_APO_ and (KCNQ1+KCNE1)_PIP2_. Only S2 and S4 were shown for clarity. Red circles highlight E160/R231 and F167/Q234 interactions in (KCNQ1+KCNE1)_APO_; and E160/R237 and F167/H240 interactions in (KCNQ1+KCNE1)_PIP2_. Arrows indicate movements of R4 (∼8 Å), the S4 loop (∼10 Å), and the S2 loop (∼6 Å). Notably, KCNE1 induced a “helix-to-loop” transition at the “S177/A178/G179, SAG” motif (red) of S2. **(C)** VCF results of KCNQ1 (black) and KCNQ1+KCNE1 (red). VCF traces recorded at +100 mV and -120 mV were normalized to compare their activation and deactivation. **(D)** The F–V relationships were fitted with a double Boltzmann equation (F_1_ and F_2_ components), and the G–V relationships were fitted with a single Boltzmann equation. **(E)** Cartoon schemes to show KCNE1 suppresses IO but enhances AO. **(F-G)** Pore radius analysis of KCNQ1_APO_, (KCNQ1+KCNE1)_APO_, and (KCNQ1+KCNE1)_PIP2_ with front and back subunits excluded for clarity. Black and orange dots indicate the residue S349 and G345. **(H)** Structural comparison of the activation gate (S349) between (KCNQ1+KCNE1)_APO_ and (KCNQ1+KCNE1)_PIP2_. **(I)** G345 in (KCNQ1+KCNE1)_PIP2_ shows larger distance (from 10.3 Å to 11.9 Å) than in KCNQ1_PIP2_ (PDB: 7XNN ^30^). **(J)** Structural comparison of selectivity filter between KCNQ1_APO_, (KCNQ1+KCNE1)_APO_, and (KCNQ1+KCNE1)_PIP2_. KCNE1 expands the T312 carbonyl oxygens from 4.4 Å to 5.6 Å. During channel opening, the G314 carbonyl oxygens undergo a ∼30° clockwise rotation, increasing their distance from 5.1 Å to 6.3 Å, and disrupting the s2 K⁺ binding site. **(K)** Representative currents of KCNQ1 and KCNQ1+KCNE1 in presence of 100 Rb^+^ or 100 K^+^ extracellular solutions. The Rb^+^/K^+^ ratio were 3.0±0.2 for KCNQ1 and 0.6±0.1 for KCNQ1+KCNE1 (p= 0.00032).

Activation of KCNQ1 VSD involves coordinated interactions between gating charges (R1-R6) on S4 and conserved residues, including E160 (E1) from S2, and the “charge transfer center” (F167 (F0) together with E170 and D202 ^44,45^). In the intermediate VSD state, E1 pairs with the second gating charge R231 (R2), and F0 interacts with Q234 (Q3); while in the activated VSD state, E1 engages with the fourth gating charge R237 (R4) and F0 interacts with H240 (H5) ^11,12,16,35^. This established KCNQ1 VSD activation pattern offers a reliable reference for validating VSD states captured in (KCNQ1+KCNE1)_APO_ and (KCNQ1+KCNE1)_PIP2_ structures. These two structures exhibited distinct VSD conformations: in (KCNQ1+KCNE1)_APO_, E1 and F0 are in close contact with R2 and Q3, respectively (**Figure 2B**), which strongly suggests that the VSD is in the intermediate state; while in (KCNQ1+KCNE1)_PIP2_, E1 and F0 are clearly pointing to R4 and H5, respectively (**Figure 2B**), indicating that its VSD is in the activated state.

Voltage-clamp fluorometry (VCF) provides a powerful tool to track VSD activation by fluorescence ^11,12,16,46^. From VCF results, the VSD activations of KCNQ1 and KCNQ1+KCNE1 both display biphasic kinetics ^11^ (*τ*_-f_ and *τ*_-s_, **Figure 2C**), with the fluorescence-voltage (F–V) relationships well-fitted by a double Boltzmann function comprising two components, F_1_ and F_2_. F_1_ corresponds to the VSD transition from the resting state to the intermediate state, and F_2_ reflects the VSD transition from the intermediate state to the activated state ^11,18,23,30^ (**Figure 2D**). However, KCNE1 significantly slows down the VSD activation, selectively promotes the resting-to-intermediate VSD transition (left-shifts F_1_ by -60 mV), and suppresses the IO state VSD-pore coupling ^16,18,23,30,36,37^ (**Figure 2D**). As a result, the VSD of KCNQ1+KCNE1 is stabilized in the intermediate state, and the channel opening is coupled to F_2_, allowing it to open exclusively in the AO state (**Figure 2D,E**).

These VCF results suggest that although KCNQ1+KCNE1 still shares the same VSD activation pattern as KCNQ1, detailed structures in the intermediate and activated states might be different. In line with this, we observed: compared to the intermediate-state VSD structures of KCNQ1 (PDB: 6MIE ^35^ and PDB: 8SIM ^34^), the VSD of (KCNQ1+KCNE1)_APO_ exhibited ∼3.5 Å displacements in S3 and S4 while maintaining S1 and S2 positions (**Figure S11**), whereas the VSD of (KCNQ1+KCNE1)_PIP2_ preserved overall S1-S4 positions, compared to the activated state of KCNQ1 (PDB: 7XNL ^30^ and PDB: 8SIK ^34^) (**Figure S11**). These KCNE1-induced VSD rearrangements together with the ∼12° rotation, likely contribute to the stabilization of the intermediate state and the markedly slower VSD activation kinetics.

Notably, in (KCNQ1+KCNE1)_APO_, KCNE1 induced a local “helix-to-loop” unfolding at the bottom “S177/A178/G179, SAG” motif of S2, extending the loop between S2 and S2-S3 linker compared to KCNQ1 alone (**Figure 2B**). As this loop mediates direct interactions with calmodulin (CaM) ^30-33,47^, this “helix-to-loop” transition is likely due to the downward movement of the cytosolic domain together with CaM (**Figure 1H**). Consistent with this, the “SAG” motif undergoes a “loop-to-helix” refolding in (KCNQ1+KCNE1)_PIP2_ open channel structure, when CaM loses interactions with the loop upon channel opening (**Figures 1G,H, 2B**).

### KCNE1-induced structural remodeling of the ion conduction pathway

We next investigated the KCNE1-induced structural remodeling of the pore domain of KCNQ1. Using HOLE program analysis ^48^, we performed pore radius analysis across KCNQ1_APO_, (KCNQ1+KCNE1)_APO_, and (KCNQ1+KCNE1)_PIP2_ structures, revealing that KCNE1 induces structural changes to the entire ion conduction pathway including selectivity filter, central cavity, and activation gate (**Figure 2F,G**). Two major structural rearrangements occurred:

First, the (KCNQ1+KCNE1)_APO_ structure reveals a closed pore, with the inner gate residue S349 blocking the ion conduction pathway (<1 Å), whereas the (KCNQ1+KCNE1)_PIP2_ structure adopts an open conformation (S349 expansion >3 Å) (**Figure 2F-H**). Notably, compared to KCNQ1_APO_, the central cavity and the activation gate of (KCNQ1+KCNE1)_APO_ exhibit horizontal expansion and a downward shift (**Figure 2G**), highlighting KCNE1’s profound structural remodeling of the pore. For instance, G345, a small-sized amino acid, forms the narrowest point of the central cavity even when the activation gate is in the open state ^19,32^ (**Figure 2G**). Previously, it was shown that G345 plays a critical role in the partial dehydration of permeating K⁺ ions, thereby regulating single-channel conductance ^19^. In (KCNQ1+KCNE1)_PIP2_, the distance between two opposite G345 residues is significantly wider than in KCNQ1_PIP2_ (PDB: 7XNN ^30^, increasing from 10.3 Å to 11.9 Å) (**Figure 2I**), which may provide a structural basis for KCNE1’s enhancement of single-channel conductance ^11,19,49^.

Second, the selectivity filter (SF) of K_V_ channels consists of a highly conserved signature sequence ("T312/I313/G314/Y315/G316, TIGYG" in KCNQ1), where backbone carbonyl oxygen atoms form four evenly distributed K⁺ ion binding sites (s1–s4) ^30-34^. Detailed structural analysis revealed that KCNE1 induces significant widening of carbonyl oxygen of T312 (4.4→5.6 Å, **Figure 2J**). During KCNQ1+KCNE1 channel opening, T312 maintains its dilated conformation, while the G314 carbonyl oxygens undergo another widening (5.1→6.4 Å, **Figure 2J**). These structural reorganizations in SF disrupt the s2 K⁺ binding site (**Figure 2J**), which strongly suggests that KCNE1 changes the function of SF.

To investigate how KCNE1 alters SF function, we performed Rb^+^/K^+^ permeability ratio and Barium (Ba^2+^, a classical potassium channel SF blocker ^15,50^) blockage experiments. For Rb⁺/K⁺ permeability ratio measurements, we analyzed tail currents at -60 mV following depolarization to +40 mV while perfusing either 100 mM Rb⁺ or 100 mM K^+^ extracellular solutions. Remarkably, while KCNQ1 alone displayed strong Rb^+^ preference (Rb^+^/K^+^ ratio = 3.0±0.2), KCNQ1+KCNE1 exhibited reversed ion selectivity (Rb^+^/K^+^ ratio = 0.6±0.1) ^16,18,51,52^ (**Figure 2K**). This changed ion preference and the disrupted s2 site in (KCNQ1+KCNE1)_PIP2_ are consistent with previous findings that Rb^+^ ions are less favored in s2 ^51,53^, which may generate an energy barrier that reduces Rb^+^ permeation. Meanwhile, the Ba^2+^ blockage experiments show that 1 mM Ba^2+^ inhibited 72±1 % of KCNQ1 currents but only 51±1 % of KCNQ1+KCNE1 currents (**Figure S12**). These functional results, consistent with our structural observations, demonstrate that KCNE1 substantially reprograms SF function.

### The intermediate-to-activated VSD transition triggers KCNQ1+KCNE1 channel opening

Combining the KCNE1-induced structural changes to both VSD and pore, our findings so far outline structural basis of how KCNE1 modulates the KCNQ1 channel: In contrast to KCNQ1 alone, which exhibits both IO and AO, the KCNQ1+KCNE1 channel remains closed when the VSD adopts the intermediate state, and the channel opening is strictly coupled to the VSD transition from the intermediate state to the activated state ^16,18,23,30,36,37^. These findings are supported by VCF results that, at 0 mV, the membrane potential under which cryo-EM structures were determined, the VSD of KCNQ1+KCNE1 predominantly occupies the intermediate state, while the pore remains mainly closed (**Figure 2D**).

### KCNE1-induced “helix-to-loop” transitions to the VSD-pore coupling

We next aimed to elucidate how KCNE1-induced structural changes affect VSD-pore coupling. A “Hand-and-Elbow” gating mechanism was proposed, in which 2 distinct groups of interactions, from the “hand site” and the “elbow site”, respectively, are responsible for the dynamic VSD-pore coupling of KCNQ1 ^11,16,18,23,30,35-37^ (**Figure 3A,B**). In this model, the S4 and the S4-S5 linker adopt a bent-arm conformation, with intra-subunit interactions at the "hand site" (between S4-S5 linker and lower S6) important for both IO and AO states, and inter-subunit interactions at the “elbow site” (between S4/S4-S5 linker and adjacent S5’/S6’) specifically responsible for the AO state (**Figure 3A,B**). Between S4 and S4-S5 linker, there is a “S4 hinge” loop that connects S4 and S4-S5 linker ^30-34^ (**Figure 3B**).

**Figure 3.**
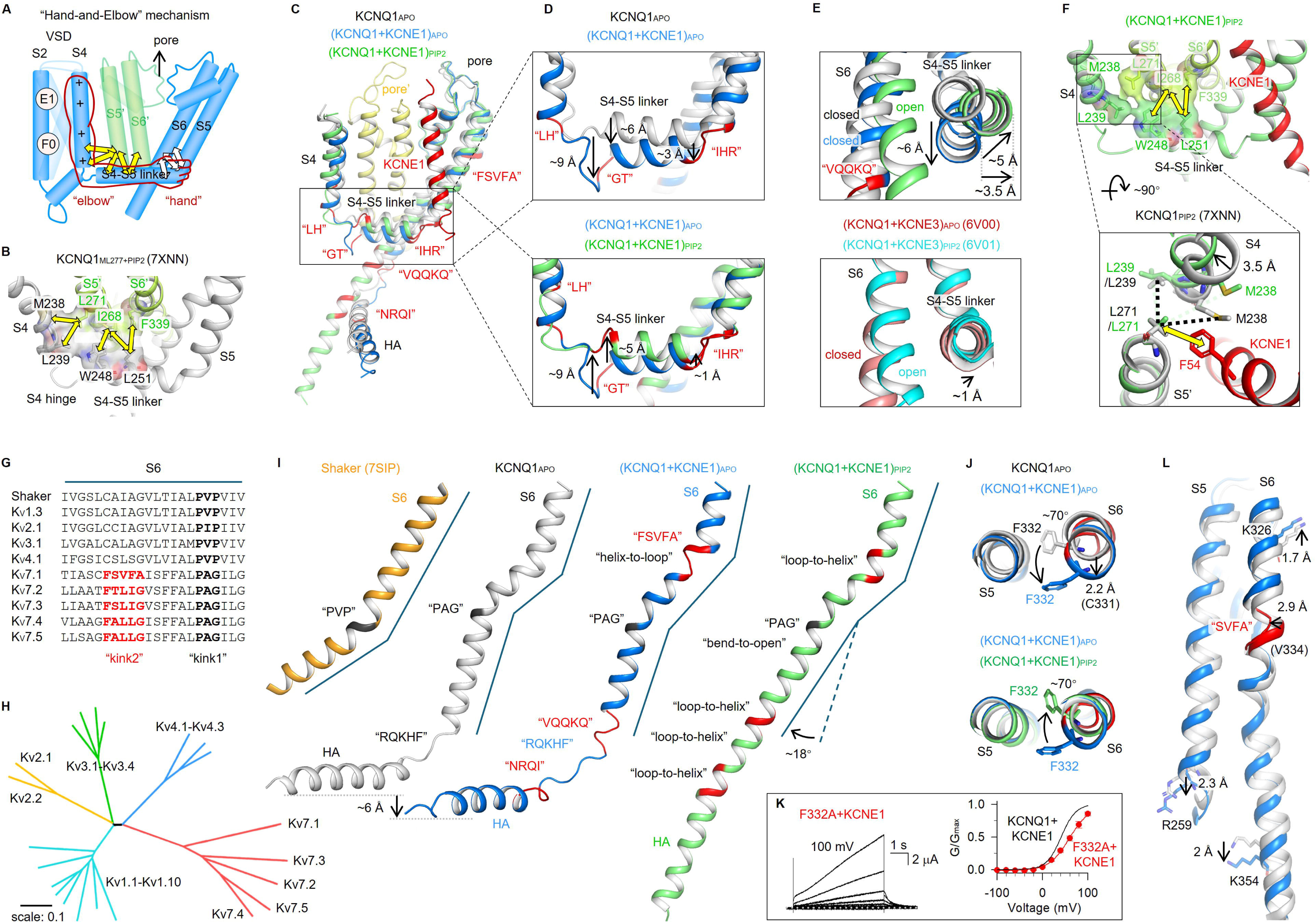
KCNE1 induced “helix↔loop” transitions at the S4-S5 linker and S6, significantly changing the VSD-pore coupling. **(A)** Cartoon scheme illustrating the “Hand-and-Elbow” gating mechanism of KCNQ1. Only two neighboring subunits (blue and green) are shown for clarity. **(B)** Five pairs of AO-specific interactions at the elbow site ^23^ were projected to KCNQ1 (PDB: 7XNN). **(C-D)** Structural comparison of KCNQ1_APO_ and (KCNQ1+KCNE1)_APO_ to show KCNE1-induced three “helix-to-loop” transitions around the S4-S5 linker: “L239/H240, LH”, “G246/T247, GT”, and “I257/H258/R259, IHR”. And these “helix-to-loop” transitions preserved during the opening of KCNQ1+KCNE1. **(E)** Structural comparison to show KCNE1-induced a ∼6 Å downward movement to the N-terminal of the S4-S5 linker. During the opening of KCNQ1+KCNE1, the S4-S5 linker undergoes a ∼5 Å upward movement (with a ∼3.5 Å horizontal expansion). The S4-S5 linker of KCNQ1+KCNE3 undergoes <1 Å upward movement with minimum horizontal expansion during opening. **(F)** KCNE1-induced remodeling to the five pairs of AO-specific interactions at the elbow site. Interactions M238/L271 and L239/L271 were abolished, but new L271/KCNE1(F54) interaction was formed. **(G)** Sequence alignment of K_V_ S6. The “kink 1” motif is conserved across all K_V_s, and “kink 2” is conserved in KCNQ channels. **(H)** Evolutionary tree analysis of all domain-swapped K_V_ channel families (K_V_1-4 and K_V_7). **(I)** KCNE1 induced three “helix-to-loop” transitions to S6: one at the “kink 2”, and two at the two ends of the S6/HA linker “RQKHF” (“VQQKQ” and “NRQI”). These loops undergo “loop-to-helix” transitions during the opening of S6, following the “bend-to-open” at “kink 1”. **(J)** Structural comparison of F332 to show KCNE1 induced a ∼70° rotation (counterclockwise) to the side chain. This rotation was restored during the channel opening. **(K)** F332A+KCNE1 currents and G–V relation (V_50_=59.3±3.0 mV). **(L)** KCNE1 induced a 2.9 Å shrink to V334, a 2.3 Å downward movement to the bottom of S5 (R259), and a 2 Å downward movement to the bottom of S6 (K354), and a 1.7 Å upward movement to the top of S6 (K326).

Remarkably, KCNE1 induced three “helix-to-loop” transitions around the S4-S5 linker: one at the “hand site” (“I257/H258/R259, IHR” motif), disturbing the direct helix connection between S4-S5 linker and S5, and two at the “elbow site” (“L239/H240, LH” and “G246/T247, GT” motifs), extending the S4 hinge from both ends. Consequently, these “helix-to-loop” transitions cause substantial downward movement of the S4-S5 linker (∼6 Å at N-terminus; ∼3 Å at C-terminus, **Figure 3C,D**). The S4 hinge dropped by an even larger distance (∼9 Å at Q244) following the VSD transition from KCNQ1_APO_’s activated state to (KCNQ1+KCNE1)_APO_’s intermediate state (**Figure 3C,D**). As an upward movement of the S4-S5 linker is required for the voltage-dependent opening, these pronounced downward movements of S4-S5 linker and S4 hinge, as well as the flexible loop connections, would likely create an energy barrier that stabilizes pore closure.

To elucidate the VSD-pore coupling that supports the KCNQ1+KCNE1 channel opening, we performed a structural comparison of (KCNQ1+KCNE1)_APO_ and (KCNQ1+KCNE1)_PIP2_. Following the VSD activation from intermediate state to activated state, the “elbow site” undergoes a large upward movement (∼9 Å for Q244, and ∼5 Å for W248), while the “hand site” moves up only ∼1 Å (at I257) (**Figure 3C,D**). Notably, from the top view, the N-terminal of S4-S5 linker undergoes a 3.5 Å horizontal expansion, driving the pore opening with a diagonal upward movement (**Figure 3E, Figure S13**). This gating motion at S4-S5 linker differs fundamentally from both KCNQ1+KCNE3 and KCNQ1, which either shows only ∼1 Å upward movement with negligible horizontal expansion (**Figure 3E**), or ∼6 Å upward movement but <1 Å horizontal expansion (**Figure S13**). These differential S4-S5 linker motions during gating provide a structural perspective that KCNQ1 and KCNQ1+KCNE3 channels favor IO, while KCNQ1+KCNE1 only opens in AO ^11,16,18,23,30,35-37^.

Using DMC analysis, we previously have identified five pairs of inter-subunit interactions at the “elbow site” that are critical for AO-specific VSD-pore coupling: M238(S4)/L271(S5’), L239(S4)/L271(S5’), W248(S4-S5 linker)/I268(S5’), L251(S4-S5 linker)/I268(S5’), and L251(S4-S5 linker)/F339(S6’) ^23^ (**Figure 3A,B**). Further investigation revealed that KCNE1 not only reshapes these interactions but also establishes new interactions between KCNQ1 and KCNE1. For example, KCNE1 displaces M238 and L239 away from L271 and abolishes interactions of M238/L271 and L239/L271 in both (KCNQ1+KCNE1)_APO_ and (KCNQ1+KCNE1)_PIP2_ (**Figure 3F**). Instead, L271 forms a new direct interaction with KCNE1-F54 (**Figure 3F**), which has been confirmed by DMC analysis (**Figure 1K,L**). Overall, these results uncover the unique VSD-pore coupling process of KCNQ1+KCNE1.

### KCNE1-induced “helix-to-loop” transitions to the S6 and HA

Unlike other K_V_ channels (e.g., the *Shaker* channel) feature a single conserved “PVP” hinge sequence (“P343/A344/G345, PAG” in KCNQ1 ^54^) in the mid-S6 to form a kink and enable a “bend-to-open” process (**Figure 3G,I**), KCNQ channels (KCNQ1-5) exhibit a distinctive dual-hinge architecture with an additional kink-forming sequence (“F332/S333/V334/F335/A336, FSVFA” in KCNQ1) above “PAG”, creating two kinks in S6 that diversify the gating (**Figure 3G,I**). In line with this observation, we performed an evolutionary analysis of K_V_ channels with domain-swapped architecture (K_V_1-4, and K_V_7), and found that clades of K_V_7 channels show a significantly higher degree of evolutionary divergence than K_V_1-4 (**Figure 3H**).

Structural comparison of KCNQ1_APO_ and (KCNQ1+KCNE1)_APO_ revealed that KCNE1 induces another three major conformational changes: 1) a ∼70° counterclockwise rotation of the conserved F332 side chain (**Figure 3J**); 2) as a result, the “FSVFA” motif undergoes a “helix-to-loop” unfolding (**Figure 3I**); 3) two more “helix-to-loop” transitions at both ends of the S6-HA linker (“V355/Q356/Q357/K358/Q359, VQQKQ” and “N365/R366/Q367/I368, NRQI”) that elongate the linker from “R360/Q361/K362/H363/F364, RQKHF” to “VQQKQRQKHFNRQ” (**Figure 3I**).

The F332 residue is well conserved among KCNQ channels (**Figure 3G**). Physically, it points to the S5 and undergoes a ∼10° clockwise rotation during KCNQ1 opening (top view of KCNQ1_APO_ and KCNQ1_PIP2_ (PDB: 7XNN ^30^), **Figure S14**). KCNE1 association, however, counterclockwise rotates F332 by ∼70° to the other side of S5, which expands the helix above F332 (2.2 Å for C331, due to the interaction with KCNE1-F53, **Figure 1K,L**), and breaks hydrogen bonds below F332 (shrinking V334 by 2.9 Å), inducing a “helix-to-loop” unfolding of the “FSVFA” motif (**Figure 3I,J,L**). To note that this “helix-to-loop” unfolding induced an upward movement to S6 above the “FSVFA” motif (e.g., 1.7 Å for K326, **Figure 3L**), and a downward shift of the S6 and S5 segments (2.3 Å for R259 in S5 and 2.0 Å for K354 in S6; **Figure 3L**). On top of these changes in S6, KCNE1 induced an additional two “helix-to-loop” transitions that elongate the S6-HA linker (**Figure 3I**). Following these changes, the entire cytosolic domain has a ∼6 Å downward movement (**Figures 1H, 3I**). These profound secondary structural changes may also contribute to establishing a higher energy barrier that stabilizes the channel in the closed state.

During the opening of KCNQ1+KCNE1, F332 rotates backward ∼70° (clockwise, **Figure 3J**). Meanwhile, the “FSVFA” motif and the “VQQKQRQKHFNRQ” loop refold to helical conformations, enabling S6 and HA to form a continuous helix, and the cytosolic domains HA/HB together with CaM undergo ∼180° rotation (**Figure 3I**). Concurrently, the “PAG” kink mediates a “bend-to-open” transition in S6 that facilitates channel opening (**Figure 3I**). Furthermore, we found F332A+KCNE1 showed a significantly right-shifted G*−*V relation (V_50_= 62.6±3.7 mV, **Figure 3K**), which confirms the importance of F332 rotation during KCNQ1+KCNE1 gating.

### A unique dual-PIP2 modulation mechanism of KCNQ1+KCNE1

The membrane phospholipid PIP2 is essential for both KCNQ1 and KCNQ1+KCNE1 channels’ gating, particularly in mediating VSD-pore coupling ^55^. Although structural studies have identified a canonical PIP2-binding site, involving basic residues from the S2-S3 linker, S4-S5 linker, and CaM ^30,32^, several key questions remain unresolved: 1) Current structures (both KCNQ1 and KCNQ1+KCNE3) show only one PIP2 binding per subunit, and PIP2 binding appears strictly limited to the activated VSD state ^30-34^, leaving it unclear whether additional PIP2 molecules participate in channel gating and whether PIP2 binding can occur in VSD other than the activated state. 2) KCNE1 enhances PIP2 sensitivity of KCNQ1 (∼100-fold) ^16,36,56^. However, the structural basis for this dramatically enhanced PIP2 sensitivity remains unknown.

Resolution of the (KCNQ1+KCNE1)_APO_ structure, prepared without adding exogenous PIP2, unexpectedly revealed well-defined PIP2 density at the canonical site (**Figure 4A**). We attribute this to endogenous PIP2 retention that survives protein purification: 1) The KCNE1 induces ∼12° counterclockwise VSD rotation, along with 6-8 Å displacements in the S2-S3 linker, S4-S5 linker, and the entire cytosolic domain may better “clamp” the head group of PIP2 (**Figure 4A**); 2) The extended S4 hinge undergoes a pronounced downward shift, positioning it closer to the neck and tail regions of PIP2 (from 6.9 Å to 4.6 Å, **Figure 4B**), which stabilizes PIP2 binding. These results not only elucidate KCNE1’s role in boosting PIP2 sensitivity and preserving endogenous PIP2 during protein preparation but also demonstrate that (KCNQ1+KCNE1)_APO_ adopts a well-coupled structure that binds PIP2 in the intermediate VSD state.

**Figure 4.**
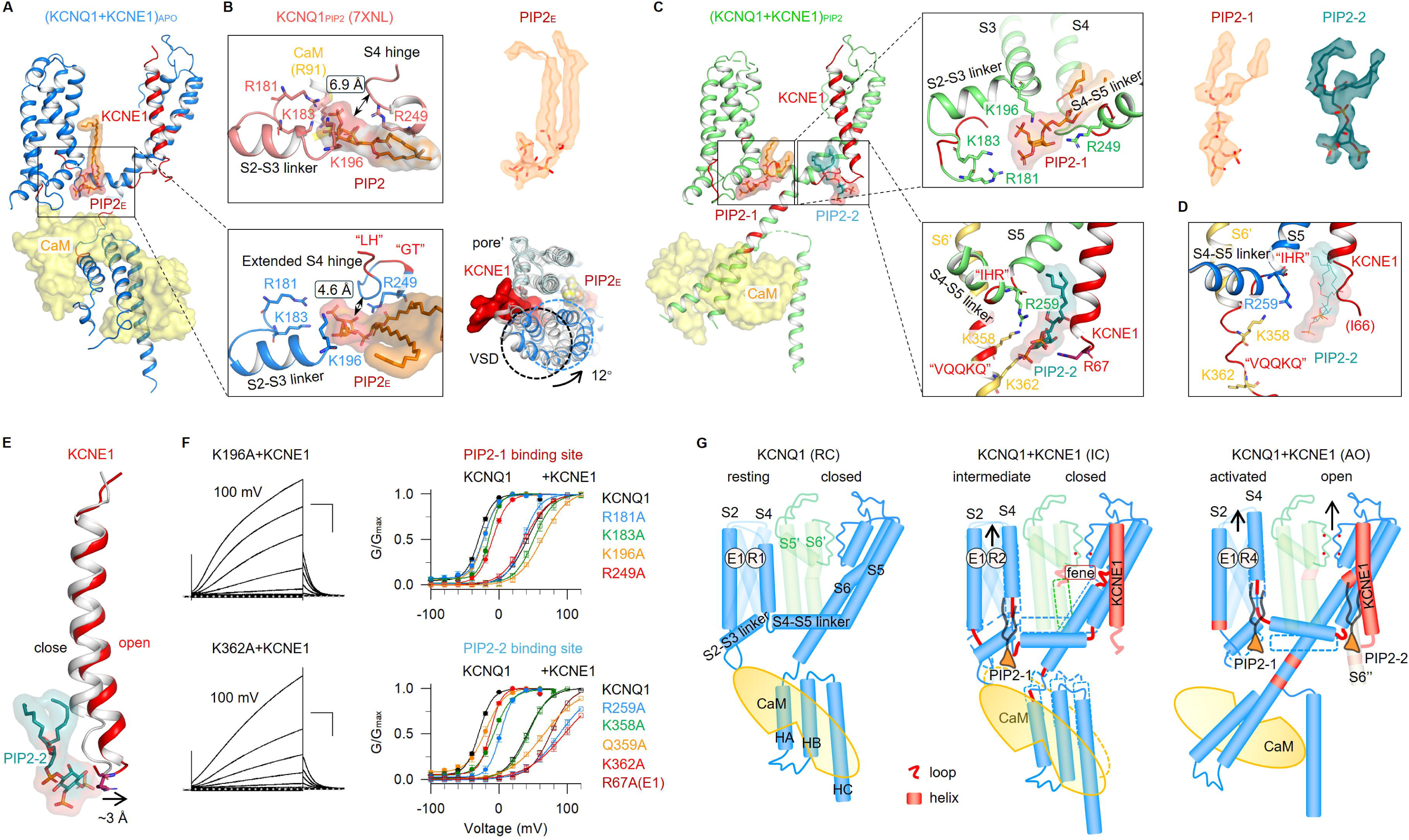
A unique dual-PIP2 modulation mechanism of KCNQ1+KCNE1. **(A)** In (KCNQ1+KCNE1)_APO_, an endogenous PIP2_E_ molecule was observed in the canonical PIP2 site (involving residues R181, K183, K196, and R249), while the CaM is in the “attached” mode. Density maps and models of PIP2_E_ were shown in orange. KCNE1-induced ∼12° VSD rotation may enhance the PIP2 sensitivity. **(B)** The extended S4 hinge in (KCNQ1+KCNE1)_APO_ shows smaller distance to the PIP2 molecule than that of in KCNQ1_PIP2_ (from 4.6 Å to 6.9 Å). **(C)** In (KCNQ1+KCNE1)_PIP2_, two PIP2 molecules (PIP2-1 and PIP2-2) were observed in each KCNQ1 subunit: PIP2-1 was in the canonical site, and PIP2-2 was binding between S4-S5 linker, S6’, and KCNE1 (involving residues R259, K358, K362, and KCNE1-R67), while the CaM is in the “detached” mode. Density maps and models of PIP2-1 and PIP2-2 were shown in orange and dark green. **(D)** Residues involved in PIP2-2 binding (R259, K358, K362, and KCNE1-R67) fall apart in the closed channel structure (KCNQ1+KCNE1)_APO_. PIP2-2 from (KCNQ1+KCNE1)_PIP2_ was also shown. **(E)** PIP2-2 binding induced ∼3 Å movement to the bottom part of KCNE1 without altering the top part. **(F)** G–V relations of Alanine mutagenesis scanning of PIP2-1 and PIP2-2 binding residues in the absence and in the presence of KCNE1. For R67 on KCNE1, KCNQ1+R67A was tested. **(G)** Cartoon schemes to show KCNE1-induced global structural remodeling to KCNQ1, and the gating process of KCNQ1+KCNE1. Briefly, KCNE1 stabilizes the VSD in the intermediate state (with a PIP2 binds to the canonical site), and induces 7 “helix-to-loop” transitions (red lines) per KCNQ1 subunit, keeping the KCNQ1+KCNE1 channel in the IC state. Upon depolarization, the VSD moves to the activated state, three loops from S6 undergo “loop-to-helix” transitions to form a continues helix along S6 and HA, and a second PIP2 binds to stabilize the KCNQ1+KCNE1 channel in the AO state.

To uncover the PIP2 modulation mechanism, we next prepared (KCNQ1+KCNE1)_EM_ samples in the presence of exogenous PIP2. Interestingly, in the (KCNQ1+KCNE1)_PIP2_ structure, we identified two distinct PIP2 molecules (PIP2-1 and PIP2-2) per subunit: PIP2-1 occupies the canonical site, coordinated by R181, K183, K196, and R249 (**Figure 4C**); while PIP2-2 binds in a new dynamic pocket formed by the S4-S5 linker (R259), adjacent S6’ (K358 and K362), and the C-terminal of KCNE1 (R67) (**Figure 4C**). These two PIP2 molecules together trigger the large cytosolic domain structural rearrangement, leading to channel opening (**Figure 4A-C**). While PIP2-1 remains stably bound throughout the gating process, PIP2-2 exhibits AO state-dependent binding, where involved binding residues move away from each other in (KCNQ1+KCNE1)_APO_ (**Figure 4D**). Notably, the PIP2-2 binding stabilizes the lower region of KCNE1, enabling clear resolution of residues R67 (directly participates in the PIP2-2 binding) and S68 in (KCNQ1+KCNE1)_PIP2_, which was missing in (KCNQ1+KCNE1)_APO_ (**Figure 4C,D**). Within the S6’, PIP2-2 interacts with residues K358 and K362, a region that undergoes the characteristic “loop-to-helix” transition during channel opening (**Figure 4C**). Furthermore, PIP2-2 binding also triggers two distinct structural rearrangements in KCNE1: 1) a “loop-to-helix” transition at the I61/M62/L63 (IML) motif, and 2) an approximately 3 Å displacement of KCNE1’s lower region without altering its upper portion (**Figure 4E**). These observations collectively highlight the dynamic nature of PIP2-2 binding and the essential mechanistic role of “helix↔loop” transitions during channel gating.

To validate the PIP2-1 and PIP2-2 binding sites, we performed Alanine scanning mutagenesis on key interacting residues. Mutations in PIP2-1 binding residues caused moderate right-shifts in G*−*V relations (ΔV ranging 7.7*−*20.2 mV, **Figure 4F**), which is consistent with previous studies ^30,55^. When co-expressed with KCNE1, the mutant-I_Ks_ channels show similarly right-shifted G*−*V relations (ΔV ranging -5.7*−*22.5 mV, **Figure 4F**). In contrast, PIP2-2 mutations produced more pronounced right-shifts (ΔV ranging 10.3*−*31.7 mV for KCNQ1 alone; and ΔV ranging 2.3*−*53.9 mV when co-expressed with KCNE1, **Figure 4F**). These findings confirm the key roles of both PIP2 in channel activation: PIP2-1 maintains stable binding at the canonical site throughout channel gating, while PIP2-2 exhibits dynamic binding in the pore and KCNE1, playing an essential role in facilitating pore opening.

In summary, our study deciphers how the KCNE1-induced global structural remodeling is coupled to the comprehensive functional modulations of KCNQ1 (**Figure 4G**). KCNE1 occupies a strategic position at the interface of three KCNQ1 subunits, inducing: 1) a ∼12° counterclockwise VSD rotation that stabilizes the intermediate state and PIP2-1 binding; 2) three “helix-to-loop” transitions around the S4-S5 linker that lower both sides of the S4-S5 linker; 3) another three “helix-to-loop” transitions in S6 and HA that shift the pore and cytosolic domains downward, stabilizing the closed conformation in the pore. Notably, a fenestration (fene, **Figure 4G**) site is induced by these secondary structure transition, which will be investigated in the following session. During KCNQ1+KCNE1 activation, VSD transition to the activated state diagonally elevates the S4-S5 linker, creating a second PIP2 binding site that bridges the S4-S5 linker, KCNE1, and S6’. This process triggers three “loop-to-helix” transitions in S6 and HA, forming a continuous helix, while the PAG kink undergoes the “bend-to-open” motion to open the channel in AO state.

### The “helix**↔**loop” transitions create a fenestration in KCNQ1+KCNE1

KCNE3, another epithelial-specific auxiliary subunit, enables the KCNQ1+KCNE3 channel to remain constitutively open across physiological voltages, helping maintain the resting membrane potential required for Cl⁻ secretion in the gut ^8,10,13,35,38,39^ (**Figure 5A**). Our prior work demonstrated that KCNE3 markedly stabilizes the IO state, resulting in a strongly left-shifted G-V relationship ^35^ (**Figure 5B,C**). These “opposite effects” of KCNE1 and KCNE3 on KCNQ1 strongly suggest that the KCNQ1+KCNE1 and KCNQ1+KCNE3 channels have fundamentally distinct structures.

**Figure 5.**
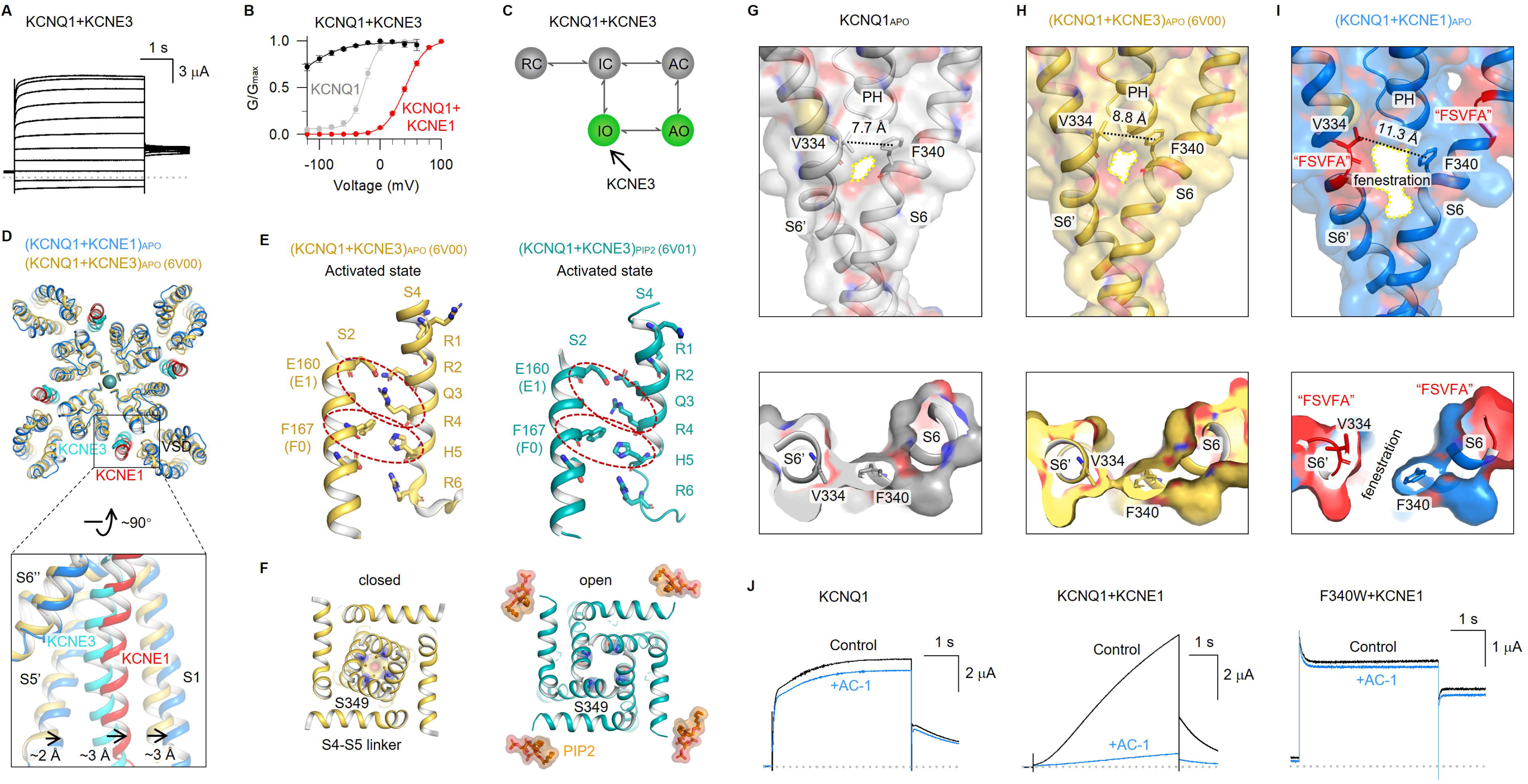
The KCNE1-induced “helix↔loop” transitions create a fenestration in the I_Ks_ channel. **(A-B)** Activation currents and G-V relation of KCNQ1+KCNE3. G-V relations of KCNQ1 (gray) and KCNQ1+KCNE1 (red) are also shown. **(C)** Cartoon schemes to show KCNE3 enhances IO. **(D)** (KCNQ1+KCNE1)_APO_ and (KCNQ1+KCNE3)_APO_ (PDB: 6V00 ^32^) comparison to show that KCNE1 induced further ∼3 Å movements to KCNE3 and S1. **(E)** Both the VSDs of (KCNQ1+KCNE3)_APO_ (PDB: 6V00) and (KCNQ1+KCNE2)_PIP2_ (PDB: 6V01) ^32^) are in activated state. Red circles highlight E160/R237 and F167/H240 interactions. **(F)** Structural comparison of the activation gate (S349) between (KCNQ1+KCNE3)_APO_ (PDB: 6V00) and (KCNQ1+KCNE2)_PIP2_ (PDB: 6V01) ^32^). **(G)** In KCNQ1_APO_, adjacent S6 segments remain tightly packed (V334↔F340 distance is 7.7 Å). **(H)** In (KCNQ1+KCNE3)_APO_, no "helix-to-loop" transition was observed, and adjacent S6 segments remain tightly packed (V334↔F340 distance is 8.8 Å). **(I)** In (KCNQ1+KCNE1)_APO_, KCNE1-induced "helix-to-loop" transitions at the "FSVFA" motifs force S6 helices to separate, creating a fenestration pocket (V334↔F340 distance is 11.3 Å). **(J)** Representative activation currents of KCNQ1, KCNQ1+KCNE1, and F340W+KCNE1 before and after adding 1 *μ*M AC-1.

We next compared the structures of the KCNQ1+KCNE1 and KCNQ1+KCNE3 channels, identifying three major differences: 1) VSD displacement: the KCNE1-induced movement of the VSD is more pronounced than that observed with KCNE3, with the VSD (S1) exhibiting an additional ∼3 Å displacement compared to the KCNQ1+KCNE3 structure (PDB: 6V00 ^32^, **Figure 5D**); 2) VSD activation state: While (KCNQ1+KCNE1)_APO_ and (KCNQ1+KCNE1)_PIP2_ exhibit intermediate and activated VSD states, respectively (**Figure 2B**), both (KCNQ1+KCNE3)_APO_ and (KCNQ1+KCNE3)_PIP2_ structures show E160/R237 and F167/H240 interactions, stabilizing the VSDs in the activated state ^32^ (**Figure 5E**); 3) PIP2 binding mechanism: unlike KCNQ1+KCNE1 utilizes a dual-PIP2 modulation mechanism (**Figure 4**), KCNQ1+KCNE3 relies on a single PIP2 molecule bound at the canonical site for channel opening (**Figure 5F**). Collectively, our structural analyses provide high-resolution insights into the distinct regulatory mechanisms of KCNE1 and KCNE3.

KCNE1-induced global secondary structural transitions also reshape the pharmacology of KCNQ1 and KCNQ1+KCNE1 channels. In the KCNQ1_APO_ and (KCNQ1+KCNE3)_APO_ (PDB: 6V00 ^32^) structures, adjacent S6 segments remain tightly packed, sealing the central cavity (V334↔F340 distances are 7.7 Å and 8.8 Å, respectively. **Figure 5G,H**). However, in (KCNQ1+KCNE1)_APO_, the KCNE1-induced "helix-to-loop" transition at the "FSVFA" motif forces S6 helices to separate, creating distinct fenestration pockets (V334↔F340 distance is 11.3 Å, **Figure 5I**). This KCNE1-dependent fenestration provides a highly selective pocket to develop KCNQ1+KCNE1 modulators. Consistent with this observation, the Seebohm lab previously has identified AC-1, a highly selective and potent blocker of KCNQ1+KCNE1 with no activity against KCNQ1 alone ^29^ (**Figure 5J**). Their functional studies demonstrated that KCNE1 binding opens fenestrations between S6 helices, enabling AC-1 binding ^29^. Our structural analysis of (KCNQ1+KCNE1)_APO_ identifies F340 as a critical residue lining the fenestration pocket (**Figure 5I**). Supporting this, we found that the F340W+KCNE1 mutant abolishes AC-1 sensitivity (**Figure 5J**), indicating that the bulky tryptophan side chain sterically blocks the fenestration, preventing AC-1 binding.

## Discussion

The distinctly slow activation of the KCNQ1+KCNE1 channel is a defining feature of its physiological function in cardiac repolarization. While extensive functional studies have probed how KCNE1 modulates both VSD activation and pore opening in KCNQ1 ^8,10-14,16,24,29,35,46,57-61^, our structural analysis provides mechanistic insights that unify and extend these observations: 1) KCNE1 stabilizes the VSD in the intermediate state, slowing down both the movement to the activated state (during activation) and the return to the resting state (during deactivation). 2) KCNE1 induces three "helix-to-loop" transitions around the S4-S5 linker, reducing the efficiency of VSD-pore coupling. The downward displacement of the S4-S5 linker may also raise the energy barrier, further stabilizing the intermediate VSD and closed pore. 3) KCNE1 promotes another three "helix-to-loop" transitions in S6 and HA, increasing pore flexibility and energetically disfavoring the "bend-to-open" transition required for channel opening. Consistent with these structural observations, functional data showed slower kinetics for pore opening than VSD activation ^16,46^. 4) Dual PIP2-mediated modulation, especially the newly discovered PIP2-2 that bridges S4-S5 linker, KCNE1 and S6’. These structural determinants undergo time-consuming "helix↔loop" transitions during gating, which may also lead to delayed pore opening. 5) A large structural rearrangement at the entire cytosolic domain, including the ∼180° rotation at HA/HB/CaM during the gating process, also slows the activation and deactivation processes.

Secondary structure transitions during ion channel gating have been previously identified across diverse channel families. For example, in non-domain-swapped HCN1, channel opening converts the S4-S5 linker from a short loop to an extended flexible loop ^62^; the non-domain-swapped hSlo1+β2_N_-β4 complex in the intermediate state reveals a “helix-to-loop” transition at its hinge glycine of S6 ^63^; the domain-swapped Na_V_1.8 undergoes a “helix-to-loop” transition in DIII-S6, similarly centered at the glycine hinge ^64^, and another “loop-to-helix” transition at the hinge between S4-S5 linker and S5 was induced by the M11 mutations in Na_V_1.7 ^65^; in ligand-gated TRPM8 channel, the C-terminal of S6 undergoes a “loop-to-helix” transition during channel opening ^66^. Here, we demonstrate that an auxiliary subunit of a K_V_ channel with domain-swapped architecture leverages large-scale “helix↔loop” transitions to profoundly reprogram channel function and pharmacology. The conservation of secondary structure transitions across evolutionarily distant channels, from non-domain-swapped K channels to domain-swapped K_V_, Na_V_, and ligand-gated TRP channels, strongly suggests these transitions represent a fundamental mechanism of ion channel gating. Beyond primary structure (sequence) and static secondary structure (*α*-helix, *β*-sheet, flexible loop, etc.) of ion channel proteins, these dynamic “helix↔loop” switches introduce a new layer of structural control that fine-tunes ion channel function.

The PIP2 modulation mechanism remains a central focus in studies of KCNQ channels ^30,34,55,67,68^, with functional studies identifying multiple potential binding sites beyond the canonical site, including the S4-S5/S6 C-termini and HA-HC regions ^56,67,69-73^. Unlike previous structures of KCNQ1 and KCNQ1+KCNE3 showing PIP2 binding only to activated VSD with open pore conformation ^30-32,34^, our study reveals a novel dual-PIP2 modulation mechanism specific to KCNQ1+KCNE1: Mandala and MacKinnon reported that the VSD state regulates PIP2 accessibility, with the "up" state favoring and the "down" state occluding PIP2 binding ^34^. Here, we found that PIP2 can bind at the canonical site with intermediate VSD state. Since KCNE1 strongly stabilizes the intermediate VSD state (V_50_ of F_1_ < -120 mV), the VSD rarely returns to the resting state under physiological voltages, permitting continuous PIP2-1 binding during channel gating. For the second PIP2-2 binding, all residues coordinating this PIP2 molecule are located in flexible loop regions that undergo substantial conformational rearrangements during gating. This reveals a new mode of dynamic modulation by PIP2. Therefore, our results provide a high spatial and temporal resolution in PIP2 binding and modulation of KCNQ1 channels.

Importantly, our high-resolution structures of KCNQ1+KCNE1 channels in both closed and open conformations provide essential information for structure-based drug discovery in LQTS. Our results revealed several unique and druggable pockets: 1) The fenestration site. The KCNE1-induced fenestration is absent in KCNQ1 and KCNQ1+KCNE3, providing a structural basis for developing highly specific KCNQ1+KCNE1 pharmacological modulators that may reduce side effects in non-cardiac tissues. Furthermore, this is a pore-embedded binding pocket. Compounds targeting this site would bypass dysfunctional pathogenic mutations that impair VSD activation and VSD-pore coupling. 2) The PC binding pocket. Our findings confirm the critical role of this conserved lipid-binding site. Studies from the Larsson lab and the Liin lab have shown that this pocket mediates the polyunsaturated fatty acids binding, and can be effectively modulated to rescue the dysfunction of LQT1 mutations ^25-27^. 3) The dual PIP2 site. Besides the canonical PIP2 site, we identified a new PIP2 site that can dynamically form during channel gating. Small molecules that bind at these sites may stabilize the channel in the open conformation. Studies from the Cui lab show that a PIP2 substitute rescues defective I_Ks_ currents when endogenous PIP2 is depleted ^74^. 4) The remodeled “elbow pocket”. While KCNE1 disrupts the ML277 binding site, its extensive restructuring of this VSD-pore coupling interface offers new opportunities for developing selective modulators. Together, these insights advance our understanding of the unique gating properties of the KCNQ1+KCNE1 channel and highlight several promising avenues for the development of next-generation therapeutics for long QT syndrome.

## Acknowledgments

We thank Professor Bert L. de Groot from Max-Planck Institute for Biophysical Chemistry for providing constructive comments on the manuscript. This work was supported by the Joint Funding of the Macau Science and Technology Development Fund and the Ministry of Science and Technology of the People’s Republic of China (0006/2021/AMJ to P.H.), by the National Natural Science Foundation of China (32171221 to P.H.), by the Macau Science and Technology Development Fund (002/2023/ALC, 006/2023/SKL, 0074/2022/A2 and 0098/2023/RIA2 to P.H.), and by Macau University of Science and Technology (FRG-23-030-SKL to P.H.). J.Z. was supported by the National Natural Science Foundation of China (grant no. 32271260), the CAS “Light of West China” Program (xbzg-zdsys-202005) and the Jiangxi Province Natural Science Foundation (grant no. 20224ACB206046).

## Author Contributions

P.H. and J.Z. conceived the project, designed the research and supervised the study. L.Z., X.L., S.W., X.C., Y.H., W.N., B.H., Z.Y., D.J, H.Z., F.L., C.X., Z.Z., J.Z., and P.H. performed experiments. L.Z., X.L., X.C., S.W., Z.Y., B.H., W.Z., J.Z., and P.H. analyzed data. H.Y., L.M., C.H., V.W., S.C., B.S., Z.J., W.Z., E.N., J.S. and P.H. provided key intellectual expertise and methodologies. L.Z., X.L., S.W., X.C., W.Z., J.Z., and P.H. wrote the manuscript with input from all authors.

## Disclosures

The authors declare no competing interests.

## Methods

### Constructs and mutagenesis

Overlap extension and high-fidelity PCR were used for making each KCNQ1 channel point mutation, which was confirmed by DNA sequencing. The cRNAs of WT KCNQ1 and all mutants were synthesized using the mMessage T7 polymerase kit (Applied Biosystems-Thermo Fisher Scientific) for oocyte injections. RNAs were kept in -80 °C.

For structural analysis, we generated the constructs which inserted the full-length KCNE1 sequence and a linker at the N-terminus of the truncated constructs of KCNQ1 (residues 76 to 620). EcoRI and NotI sites were used for cloning the (KCNQ1+KCNE1)_EM_ construct into the pEGBacMam expression vector with a C-terminal Maltose Binding Protein (MBP)-10xHis tag. The human CaM gene was cloned into BacMam expression vector without any tags. (KCNQ1+KCNE1)_EM_ and CaM complex were heterologously expressed in Human Embryonic Kidney (HEK) 293F cells. When cell density reached 2.0-3.0x10^6^ cells/mL, the cells were cotransfected with plasmids at mass ratio 10:1 for (KCNQ1+KCNE1)_EM_ and CaM. For 1-liter HEK 293F cell culture, the plasmids (∼1 mg) were premixed with linear polyethylenimines (PEIs) (MKbio) in 50-ml fresh medium for 15 to 30 min. The mixture was then added into cell culture followed by 15-min incubation. After 24 hours incubation at 37 °C, 10 mM sodium butyrate was added to induce the protein expression at 30 °C. Cells were harvested after 48 h, then flash-frozen in liquid nitrogen and stored at −80 °C until needed.

Cell pellets were resuspended in hypotonic buffer (20 mM Tris-HCl, pH 8.0, 20 mM KCl, 0.5 mM MgCl_2_, 2mM DTT) supplemented with a protease inhibitor cocktail (Selleck) for 40 min with gentle agitation. Crude cell membranes were collected by ultra-centrifugation at 32000 rpm for 45 min. The membranes were then re-suspended and solubilized in buffer containing 20 mM Tris-HCl, pH 8.0, 150 mM KCl, 2mM DTT, 0.5% LMNG:CHS (10:1, w/w) for 2-2.5 h at 4°C. After centrifugation at 32000 rpm for 45 min, the supernatant was incubated with amylose resin (NEB) at 4°C for 2 h. Detergent was exchanged on resin by a series of washing steps in 20 mM Tris-HCl, pH 8.0, 150 mM KCl, 2 mM DTT supplemented with different detergents: first 0.1% LMNG, 0.01% CHS, 0.1% GDN, then 0.1% GDN and 0.05% GDN for 20 column volumes each. The protein was eluted with 40 mM Maltose in wash buffer, and subsequently concentrated by a 100-kDa concentrator (Millipore) before being injected onto a Superose 6 Column (GE Healthcare) equilibrated with 20 mM Tris-HCl, pH 8.0, 150 mM KCl, 2mM DTT, 0.03% GDN. The peak fractions were pooled, concentrated to 4-5 mg/mL using a 100-kDa MWCO centrifugal device (Millipore) before cryo-EM sample preparation. For the (KCNQ1+KCNE1)_PIP2_ sample, the purified protein was incubated with 1mM PIP2 (Echelon) for 30 min.

### Cryo-EM sample preparation and data acquisition

For grids preparation, 2.5-3.0 µL of concentrated protein complex was loaded onto glow-discharged holey carbon grids (Quantifoil Au R1.2/1.3, 300 mesh) at 4 °C under 100% humidity. Grids were blotted for 3.5 s and plunge-frozen in liquid ethane using a Vitrobot Mark IV (FEI). Micrographs were acquired on a Titan Krios microscope (FEI) operating at a voltage of 300 kV. Summary of detailed data collection was shown in **Table 1**.

### Cryo-EM data processing

Images of all datasets were imported into cryoSPARC v4.1.1. After motion corrected, electron-dose weighted and CTF estimation, the initial particles were performed by cryoSPARC blob picker and processed 2D classification to generate a template. For (KCNQ1+KCNE1)_APO_ datasets, after two rounds of 2D classification, the good particles proceeded to Ab-initio reconstruction and heterogeneous refinement. Then we used the best class of particles to generate a template. And the particles were performed by cryoSPARC template picker. Auto-picked particles were visually examined to remove false positives and were further cleaned up by multiple rounds of 2D classification. The good particles proceeded to two rounds of Ab-initio reconstruction and heterogeneous refinement. The final particle sets were re-extracted with original box size and further applied for final nonuniform refinement and local refinement. For (KCNQ1+KCNE1)_APO_ datasets, we merged the particles of the two best classes to remove duplicates and the final particle sets were re-extracted with original box size and further applied for final nonuniform refinement and local refinement.

### Model building and refinement for cryo-EM structures

Auto-sharpen of the Phenix program was used for map sharpening. The reference models, PDBs: 6UZZ, 6V00, and 6V01 were rigid-body fit into the EM density map using ChimeraX. The fit was further adjusted using the jiggle fit function in Coot. Further manual adjustment with the real-space refine zone function in Coot was used to generate an atomic model. The generated model was further refined using the real_space_refine tool in Phenix. MolProbity and Mtriage were used for validation. The pore radii was calculated using HOLE. PyMOL3.1.3 and ChimeraX were used to further analyze the structure and generate figures.

### Oocyte preparation and ion channel expression

Mature oocytes (at stage V or VI) were obtained from *Xenopus laevis* by laparotomy, following the protocol approved by the Animal Studies Committee of Macau University of Science and Technology (Protocol #: MUST-NSFC-2021022601HPP). Collagenase (Sigma Aldrich) at 0.5 mg/ml concentration was used to digest oocytes. For cRNA micro-injection, WT or mutant KCNQ1 cRNAs (9.2 ng) with or without KCNE1 cRNA were injected into each oocyte with a 4:1 KCNQ1:KCNE1 weight ratio. This allows a saturate KCNE1 association to KCNQ1 ^18,75^. The KCNQ1:CiVSP weight ratio was kept at 4:1 to obtain a suitable PIP2 depletion rate. Injected cells were incubated in ND96 solution (in mM): 96 NaCl, 2 KCl, 1.8 CaCl_2_, 1 MgCl_2_, 5 HEPES, 2.5 CH_3_COCO_2_Na, 1:100 Pen-Strep, pH 7.6) at 18°C for 2-6 days for electrophysiology recordings.

### Two-electrode voltage clamp (TEVC) and voltage-clamp fluorometry (VCF)

Microelectrodes (Sutter Instrument) were made with a Sutter (P-1000) puller with 1-3 MΩ resistances when filled with 3 M KCl. The extracellular solution was ND96 solution without CH_3_COCO_2_Na. Currents were recorded with an OC-725D TEVC amplifier (Warner). Currents were sampled at 1 kHz and low-pass-filtered at 2 kHz. For VCF experiments, oocytes were incubated on ice for 30 min in labeling solution: 10 μM Alexa 488 C5-maleimide (Molecular Probes) in 100 mM K^+^ solution. After labeling, oocytes were washed three times with the ND96 solution before VCF recordings. All recordings were performed at room temperature of 20–22 °C.

### Electrophysiology Data analysis

Data was analyzed with Clampfit (Axon Instruments), Sigmaplot (SPSS), and IGOR (Wavemetrics). Due to photo-bleaching, fluorescence signals were baseline subtracted. G–V and F–V curves were fitted with single or double Boltzmann equations in the form of 1/(1+exp(−z\**F*\*(*V*−*V*_1/2_)/*RT*)), where *V* is the voltage, z is the equivalent valence, *V*_1/2_ is the half-maximal voltage, *F* is the Faraday constant, *R* is the gas constant, and *T* is the absolute temperature. For double mutant cycle analysis, the activation energy was measured by Δ*G*=*-zFV_1/2_*. *F* is the Faraday constant. Both *z* and *V_1/2_*were estimated by fitting G–V relations of each channel with a single Boltzmann equation.

### Statistical analysis

Averaged data were presented as mean ± standard error of mean (SEM) with n specifying the number of independent experiments. Statistical analyses (t-test, paired t-test, one-way ANOVA and post-hoc mean comparison Tukey test or Dunnett test) were performed with Sigmaplot (SPSS) and R software (4.1.2 version, multcomp package). Statistical significance was set as “*” for p< 0.05, “**” for p< 0.01, and “***” for p< 0.001.

## Notes

### Competing Interest Statement

The authors have declared no competing interest.

