## Supplementary File for "Secondary structure transitions and dual PIP2 binding define cardiac KCNQ1-KCNE1 channel gating"

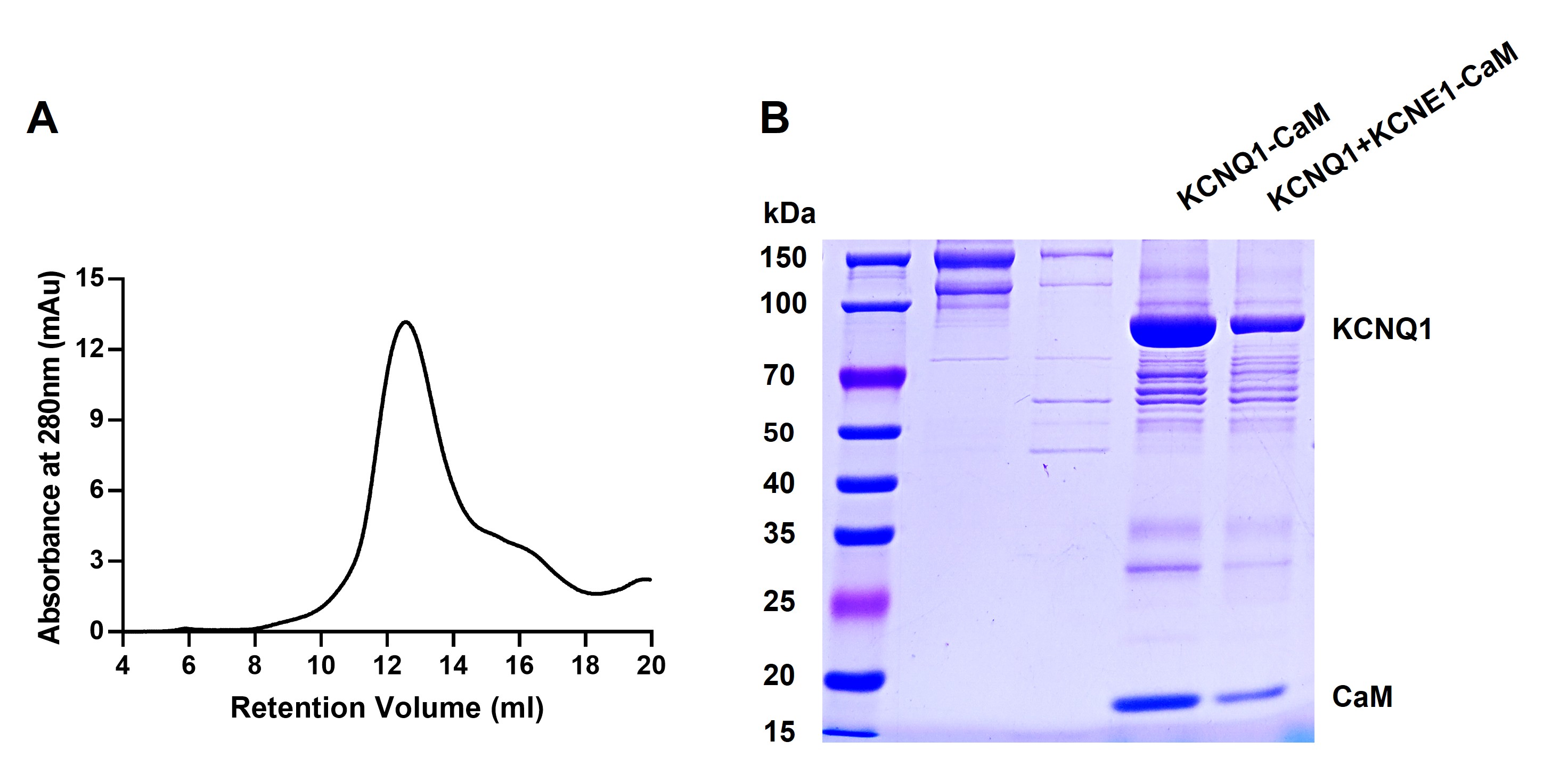
**Supplementary Figures**

**Figure S1. Initial attempts with full-length human KCNQ1 co-expressed with KCNE1 yielded only KCNQ1 protein after purification. (A)** Size-exclusion chromatography of KCNQ1+KCNE1-CaM on Superose 6 (GE Healthcare). **(B)** SDS-PAGE analysis of the KCNQ1+KCNE1-CaM and KCNQ1-CaM.

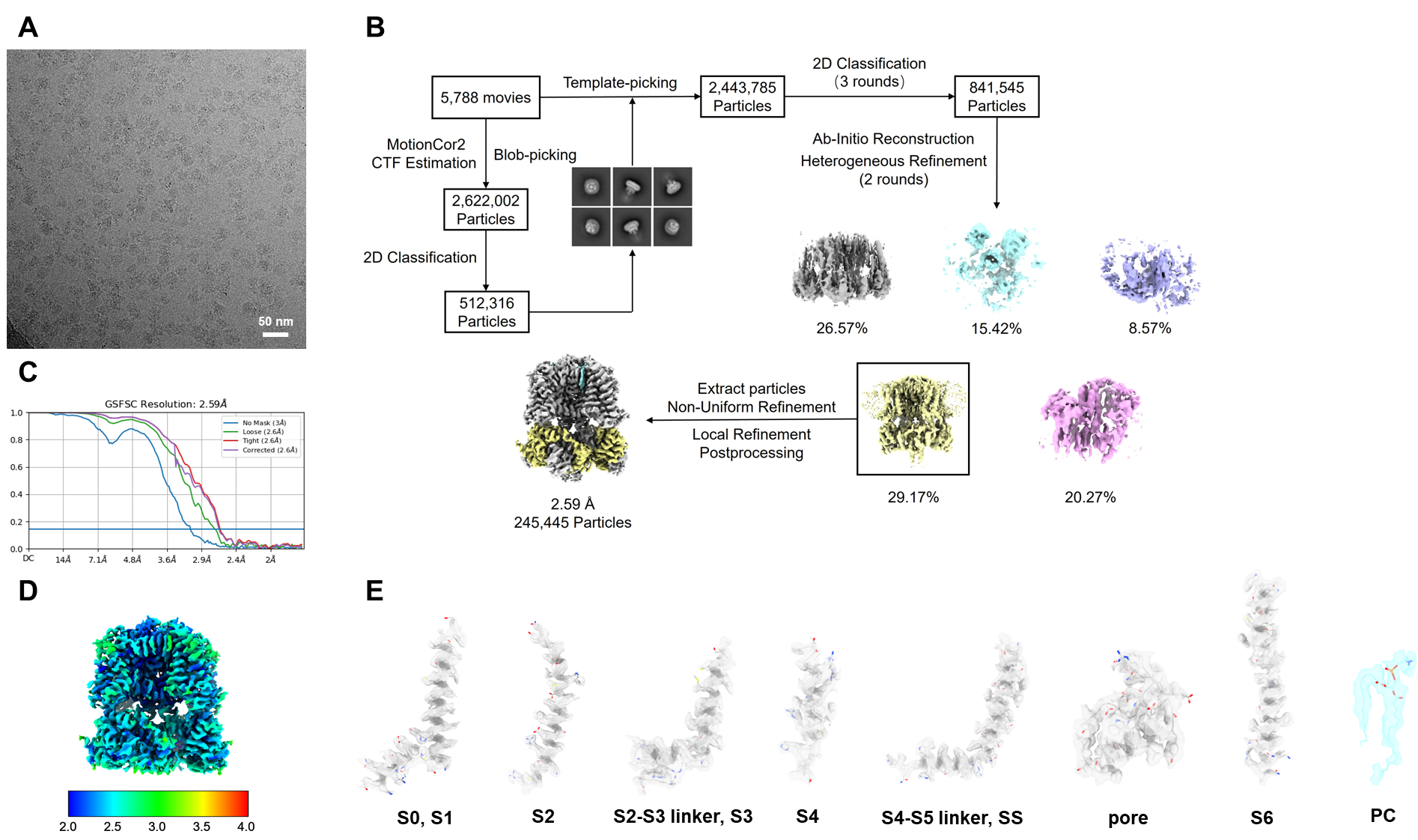
**Figure S2. Structure determination of hKCNQ1_APO_.** **(A)** A representative cryo-EM micrograph of hKCNQ1_APO_. **(B)** Flowchart of hKCNQ1_APO_ structure determination. **(C)** FSC of the final map. **(D)** Local resolution of the channel complex calculated by Blocres software. **(E)** Cryo-EM densities for various TMs in the hKCNQ1_APO_.

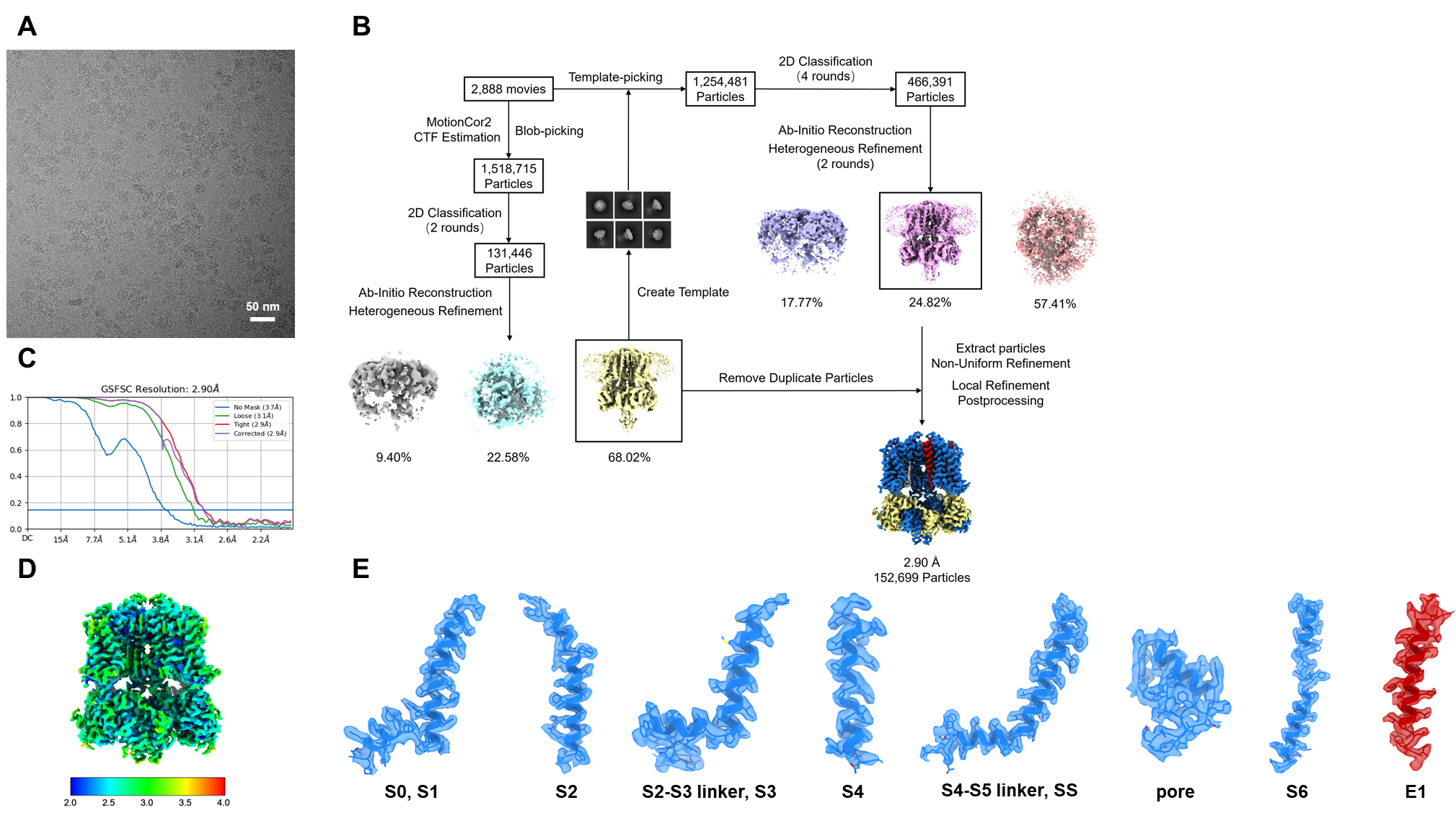
**Figure S3. Structure determination of (KCNQ1+KCNE1)_APO_. (A)** A representative cryo-EM micrograph of (KCNQ1+KCNE1)_APO_. **(B)** Flowchart of (KCNQ1+KCNE1)_APO_ structure determination. **(C)** FSC of the final map. **(D)** Local resolution of the channel complex calculated by Blocres software. **(E)** Cryo-EM densities for various TMs in the (KCNQ1+KCNE1)_APO_.

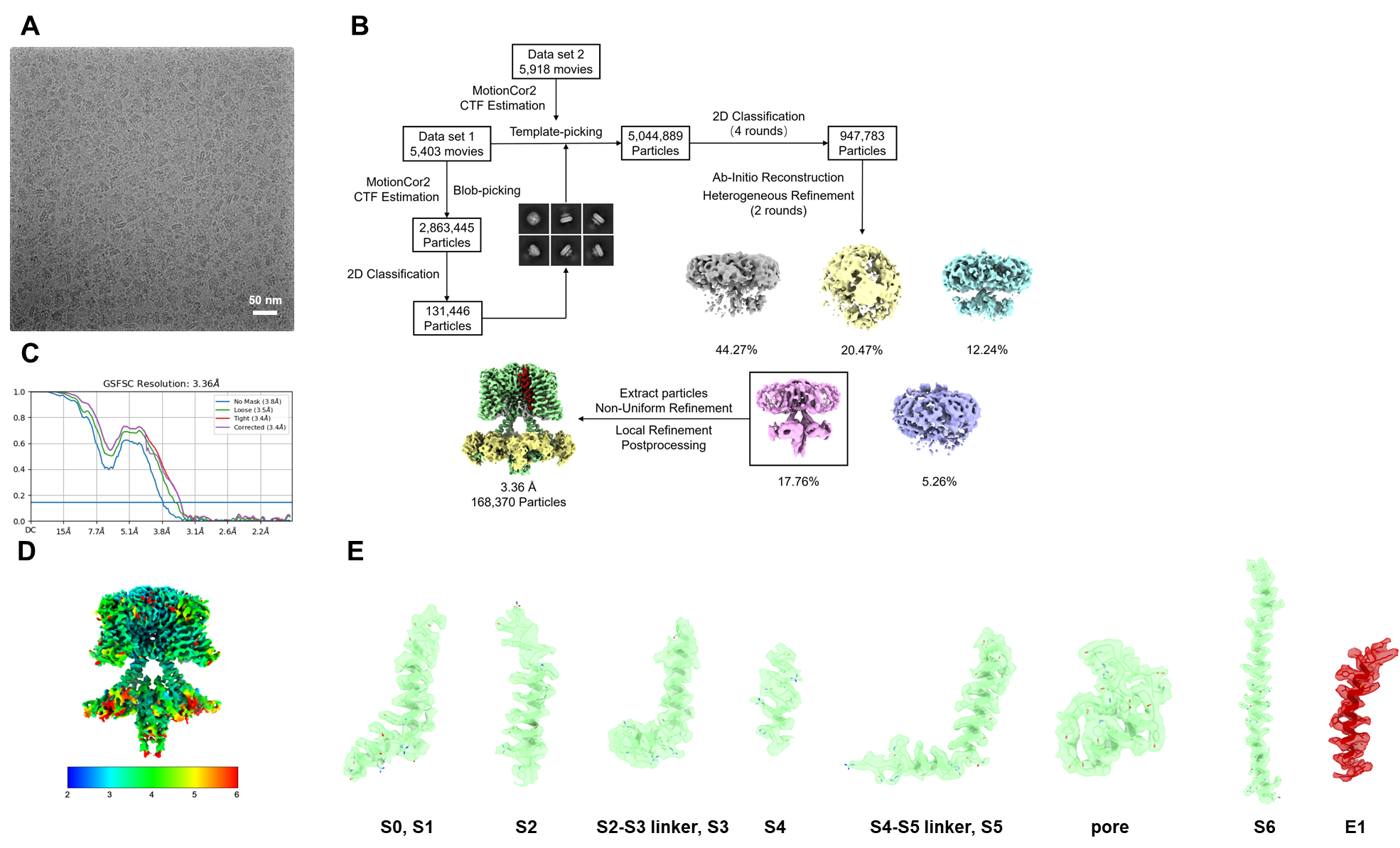
**Figure S4. Structure determination of (KCNQ1+KCNE1)_PIP2_. (A)** A representative cryo-EM micrograph of (KCNQ1+KCNE1)_PIP2_. **(B)** Flowchart of (KCNQ1+KCNE1)_PIP2_ structure determination. **(C)** FSC of the final map. **(D)** Local resolution of the channel complex calculated by Blocres software. **(E)** Cryo-EM densities for various TMs in the (KCNQ1+KCNE1)_PIP2_.

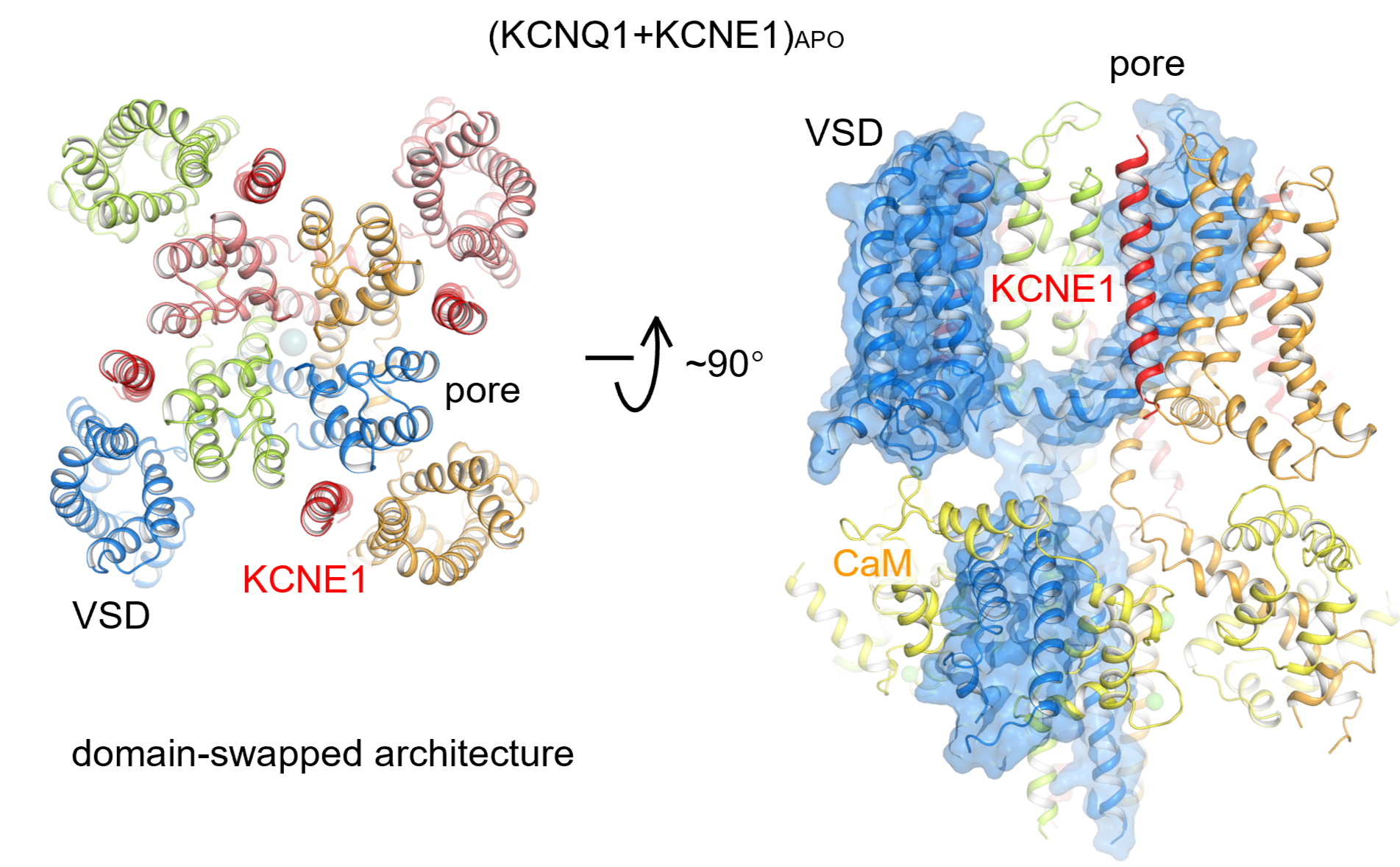

**Figure S5. The KCNQ1+KCNE1 structures still follow the domain-swapped architecture.**

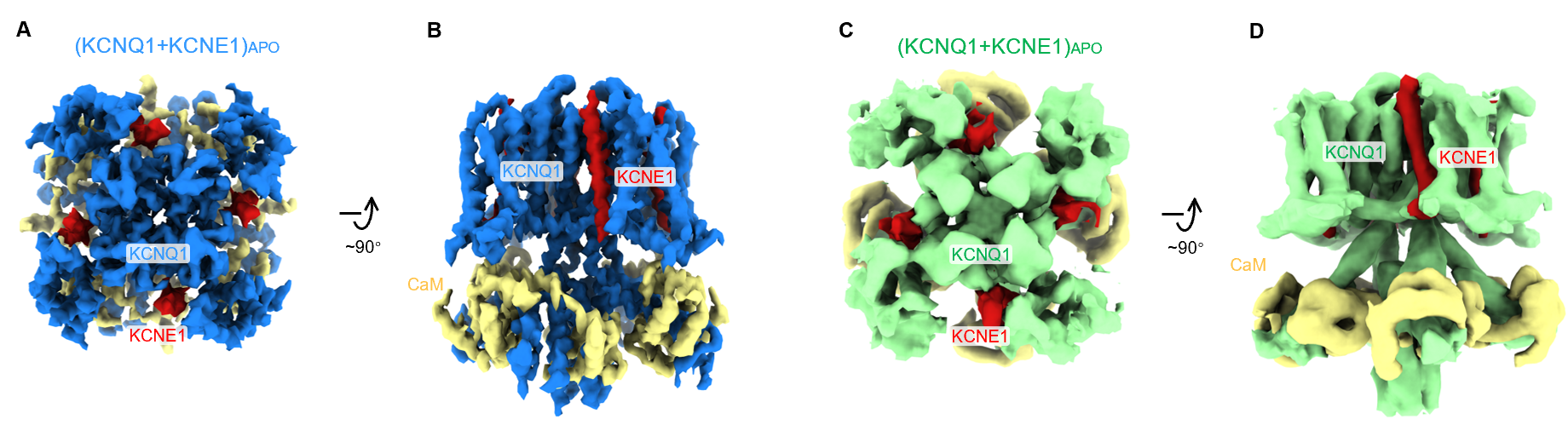

**Figure S6. Structure determinations of (KCNQ1+KCNE1)_APO_ and (KCNQ1+KCNE1)_PIP2_ in C1 symmetry**, confirming the 4:4 stoichiometry between KCNQ1 and KCNE1.

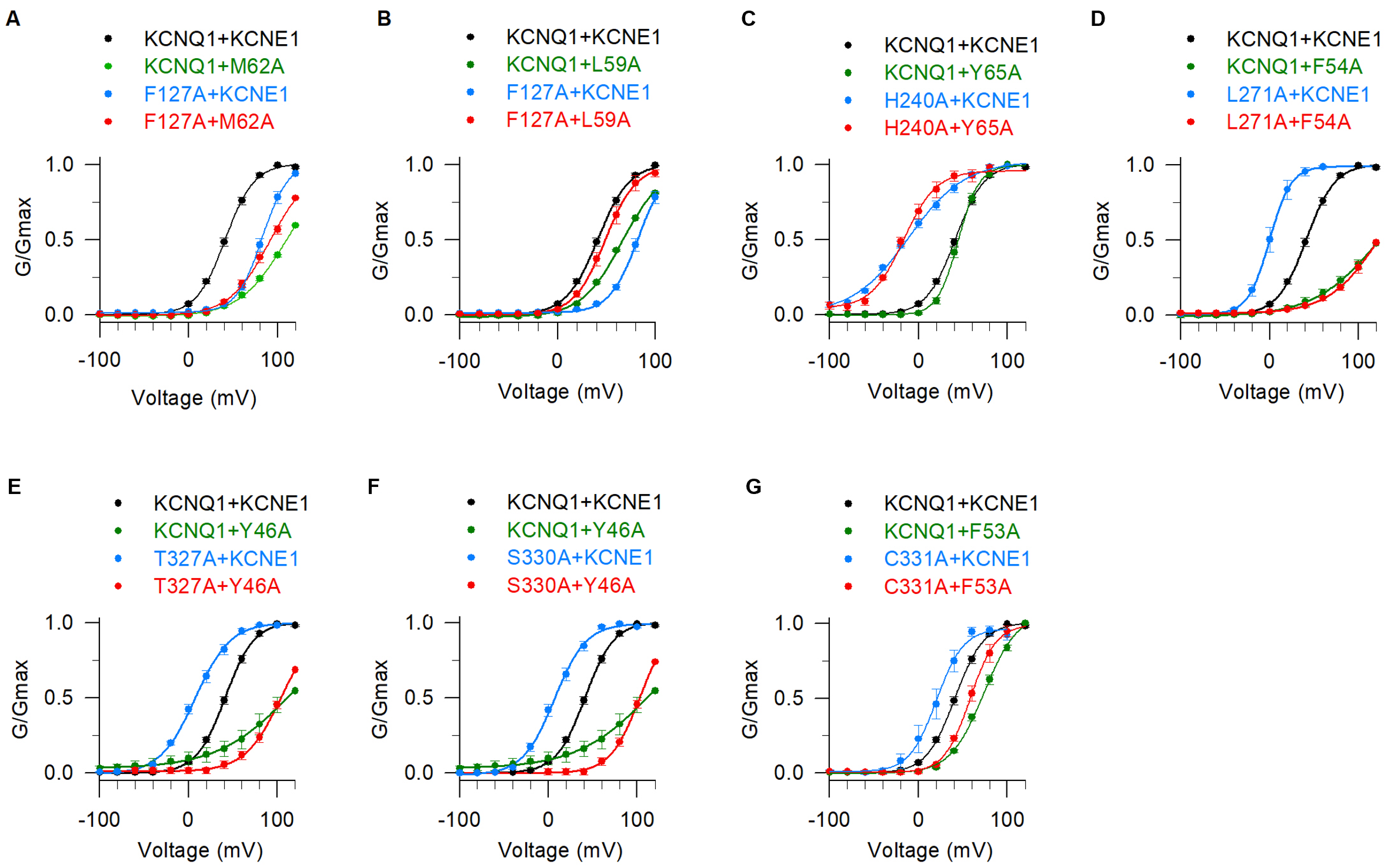

**Figure S7. DMC analysis to identify seven additional interaction pairs between KCNE1 and KCNQ1.** **(A)** Double mutant cycle analysis between KCNQ1-F127 and KCNE1-M62 with *ΔΔG* =4.7 kcal/mol. **(B)** Double mutant cycle analysis between KCNQ1-F127 and KCNE1-L59 with *ΔΔG* =4.6 kcal/mol. **(C)** Double mutant cycle analysis between KCNQ1-H240 and KCNE1-Y65 with *ΔΔG* =2.6 kcal/mol. **(D)** Double mutant cycle analysis between KCNQ1-L271 and KCNE1-F54 with *ΔΔG* =5.0 kcal/mol. **(E)** Double mutant cycle analysis between KCNQ1-T327 and KCNE1-Y46 with *ΔΔG* =8.4 kcal/mol. **(F)** Double mutant cycle analysis between KCNQ1-S330 and KCNE1-Y46 with *ΔΔG* =6.4 kcal/mol. **(G)** Double mutant cycle analysis between KCNQ1-C331 and KCNE1-F53 with *ΔΔG* =1.9 kcal/mol.

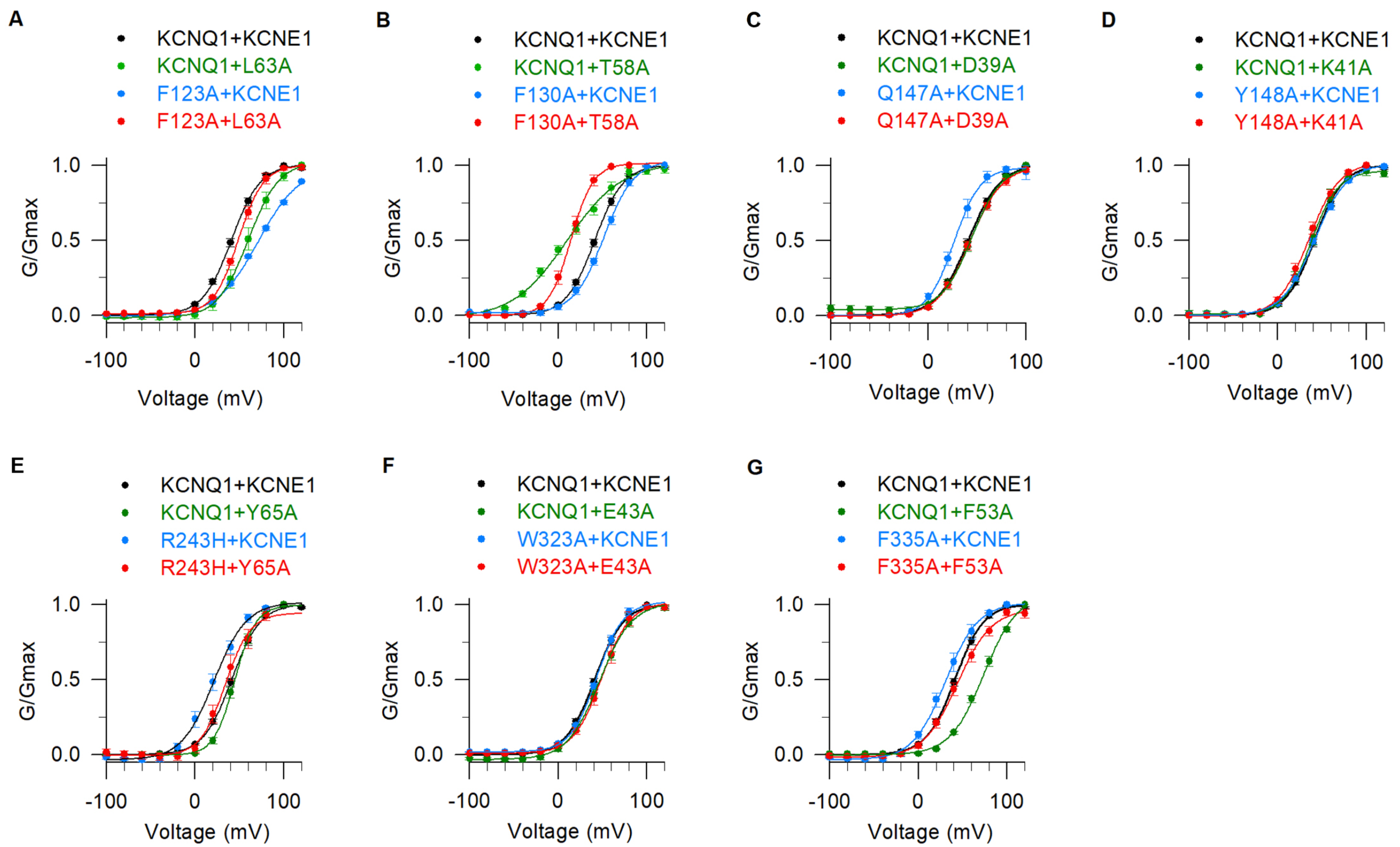

**Figure S8. DMC analysis to identify seven non-interacting residue pairs.** **(A)** Double mutant cycle analysis between KCNQ1-F123 and KCNE1-L63 with *ΔΔG* =0.41 kcal/mol. **(B)** Double mutant cycle analysis between KCNQ1-F130 and KCNE1-T58 with *ΔΔG* =0.43 kcal/mol. **(C)** Double mutant cycle analysis between KCNQ1-Q147 and KCNE1-D39 with *ΔΔG* =0.35 kcal/mol. **(D)** Double mutant cycle analysis between KCNQ1-Y148 and KCNE1-K41 with *ΔΔG* =0.01 kcal/mol. **(E)** Double mutant cycle analysis between KCNQ1-R243 and KCNE1-Y65 with *ΔΔG* =0.13 kcal/mol. **(F)** Double mutant cycle analysis between KCNQ1-W323 and KCNE1-E43 with *ΔΔG* =0.24 kcal/mol. **(G)** Double mutant cycle analysis between KCNQ1-F335 and KCNE1-F53 with *ΔΔG* =0.99 kcal/mol.

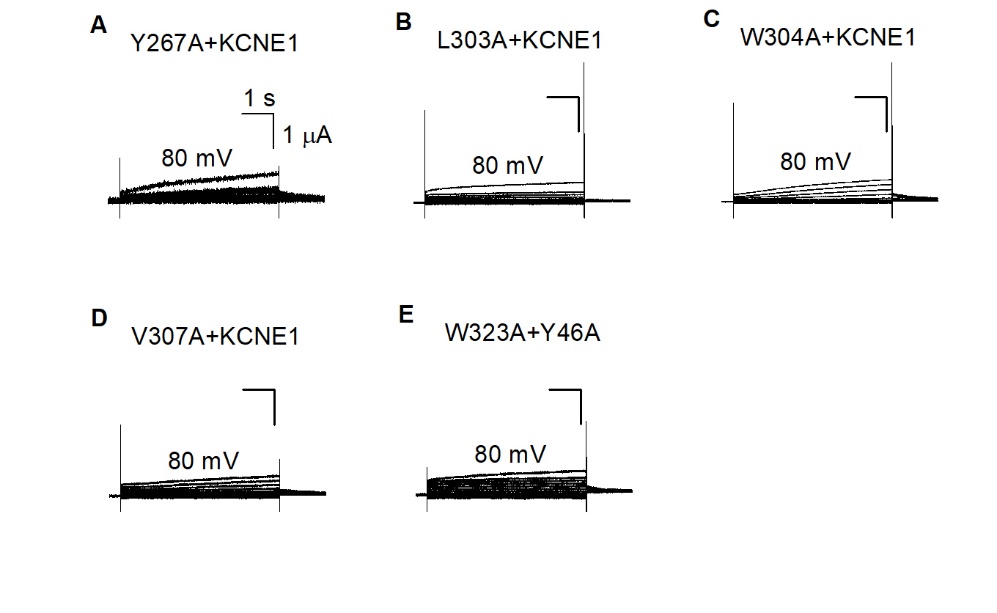

**Figure S9. KCNQ1 or KCNE1 mutations that lose the mutant-I_Ks_ current.** **(A-E)** Y267A+KCNE1, L303A+KCNE1, W304A+KCNE1, V307A+KCNE1, and W323A+Y46A show small I_Ks_ currents, which preclude DMC analysis to quantify their interactions.

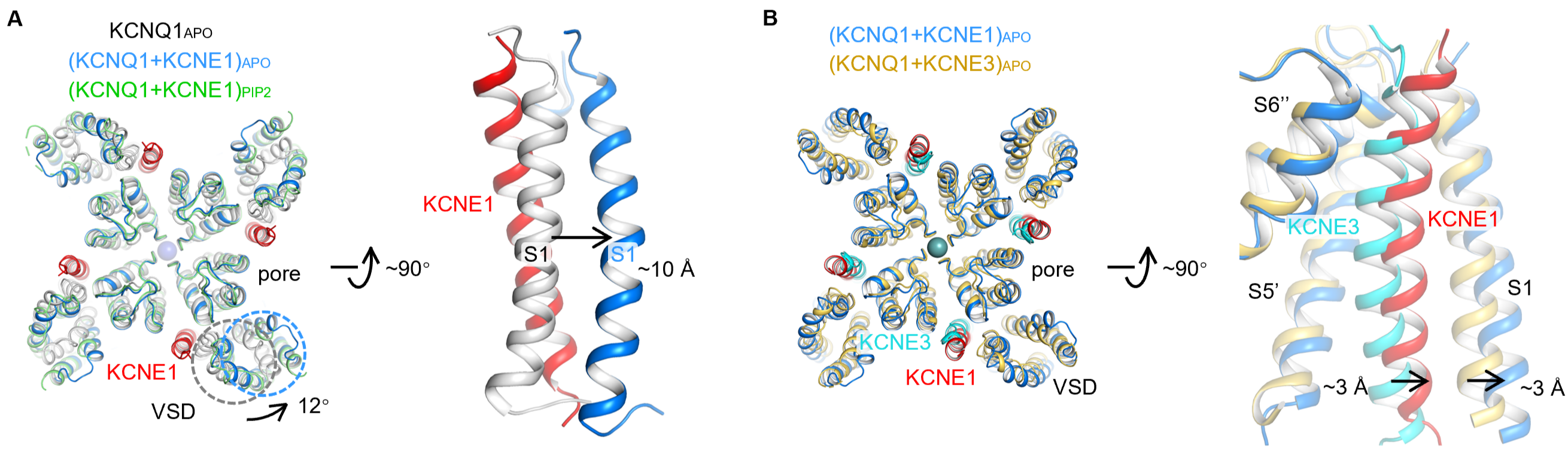

**Figure S10. KCNE1-induced conformational changes to the VSD.** Top and side views of KCNQ1_APO_, (KCNQ1+KCNE1)_APO_, and (KCNQ1+KCNE1)_PIP2_ to show KCNE1 indued a ~12° rotation to the KCNQ1 VSD (counterclockwise), and a ~10 Å movement to the S1 segment. Structures were aligned to the filter.

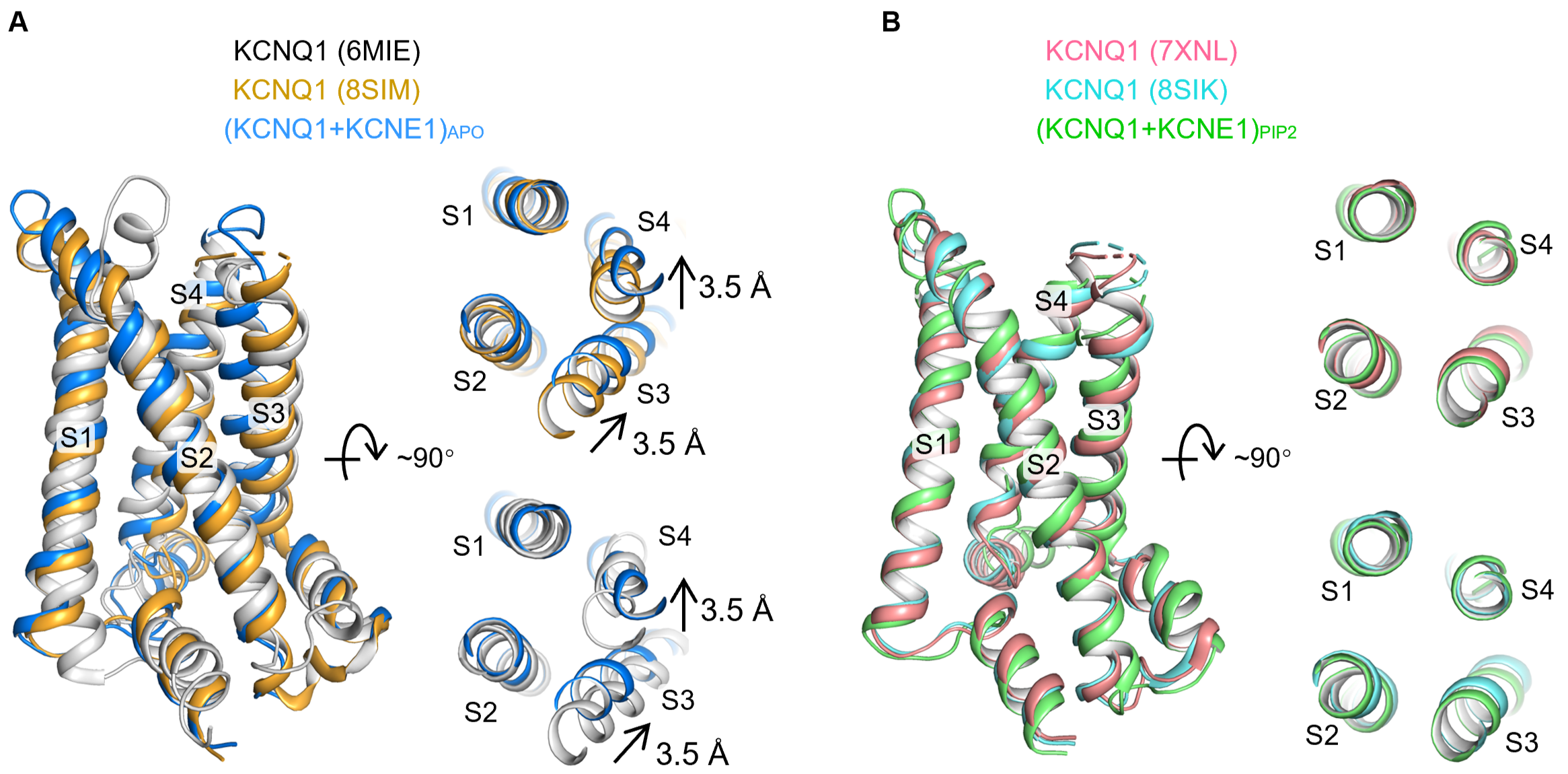

**Figure S11. KCNQ1 and KCNQ1+KCNE1 show different intermediate and activated VSD structures. (A)** Structural comparison of intermediate VSD between KCNQ1 (PDB: 6MIE ^1^ and PDB: 8SIM^2^) and (KCNQ1+KCNE1)_APO_ exhibited 3.5 Å displacements in S3-S4 while maintaining S1-S2 positions. **(B)** Structural comparison of activated VSD between KCNQ1 (PDB: 7XNL ^3^ and PDB: 8SIK ^2^) and (KCNQ1+KCNE1)_PIP2_ preserved overall S1-S4 positions.

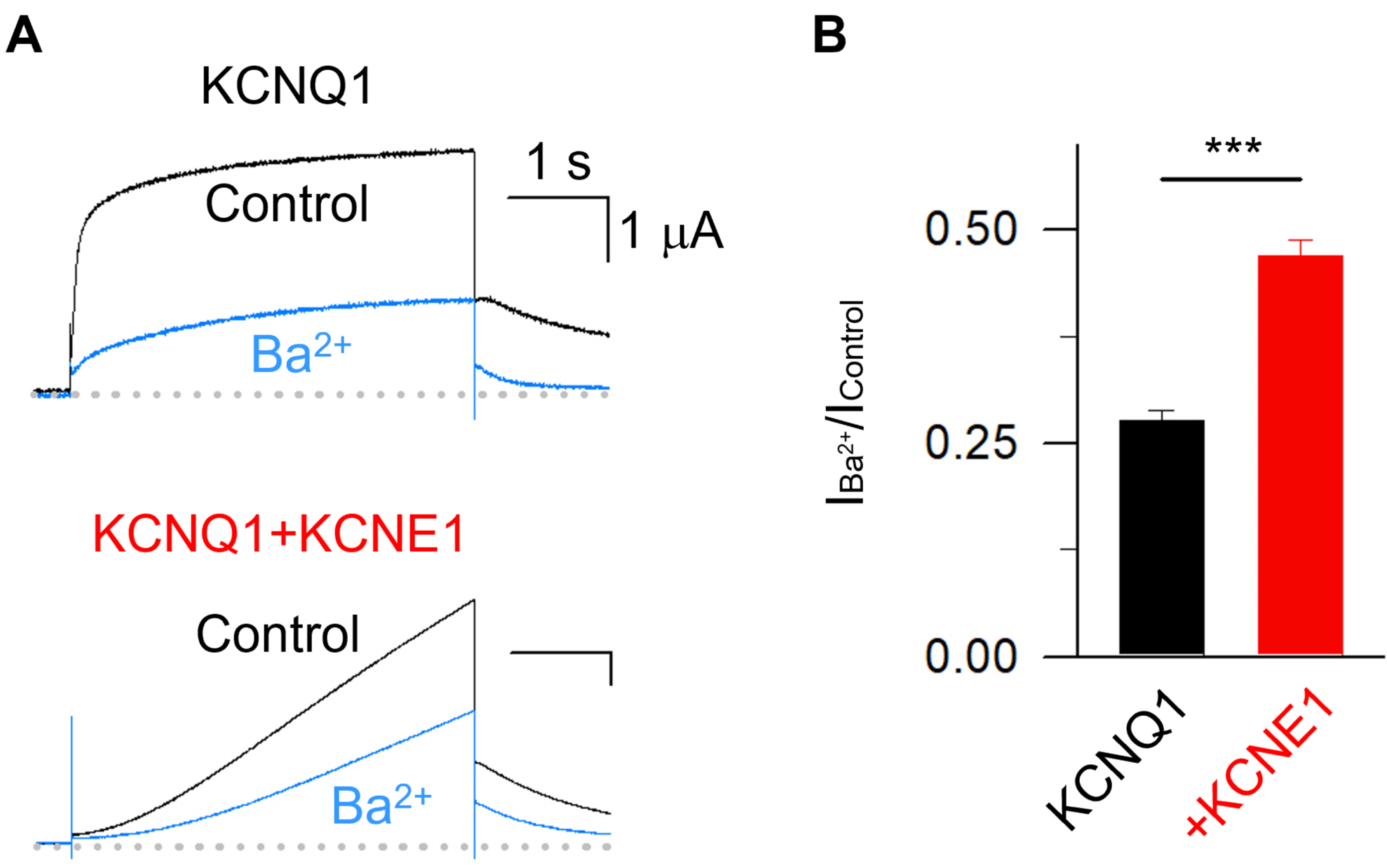

**Figure S12. Ba^2+^ blockage experiments to show KCNQ1 and KCNQ1+KCNE1 channels have different Ba^2+^ sensitivity. (A-B)** Representative currents of KCNQ1 and KCNQ1+KCNE1 before and after adding 1 mM Ba^2+^. The Ba^2+^-sensitive current (I_control-_I_Ba_^2+^)/I_control_ were 0.72±0.01 for KCNQ1 and 0.51±0.01 for KCNQ1+KCNE1.

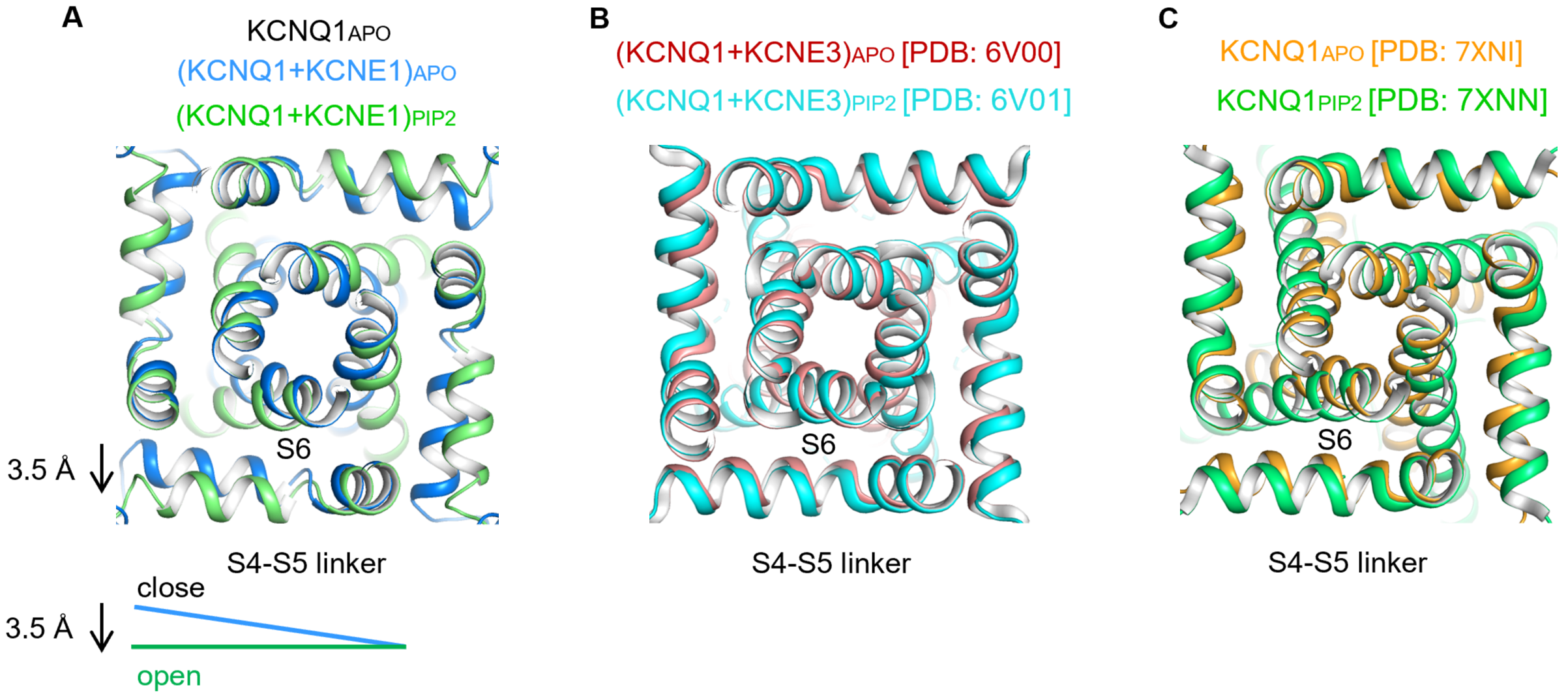

**Figure S13. KCNQ1+KCNE1 shows different S4-S5 linker motion during channel opening from KCNQ1 and KCNQ1+KCNE3 channels. (A)** Structural comparison of the activation gate (S4-S5 linker and S6) between (KCNQ1+KCNE1)_APO_ and (KCNQ1+KCNE1)_PIP2_. The S4-S5 linker shows 3.5 Å horizontal expansion during the channel opening. **(B)** Structural comparison of the activation gate between (KCNQ1+KCNE3)_APO_ (PDB: 6V00 ^4^) and (KCNQ1+KCNE3)_PIP2_ (PDB: 6V01 ^4^). The S4-S5 linker shows minimum horizontal expansion during the channel opening. **(C)** Structural comparison of the activation gate between KCNQ1_APO_ (PDB: 7XNI ^3^) and KCNQ1_PIP2_ (PDB: 7XNN ^3^). The S4-S5 linker shows <1 Å horizontal expansion during the channel opening.

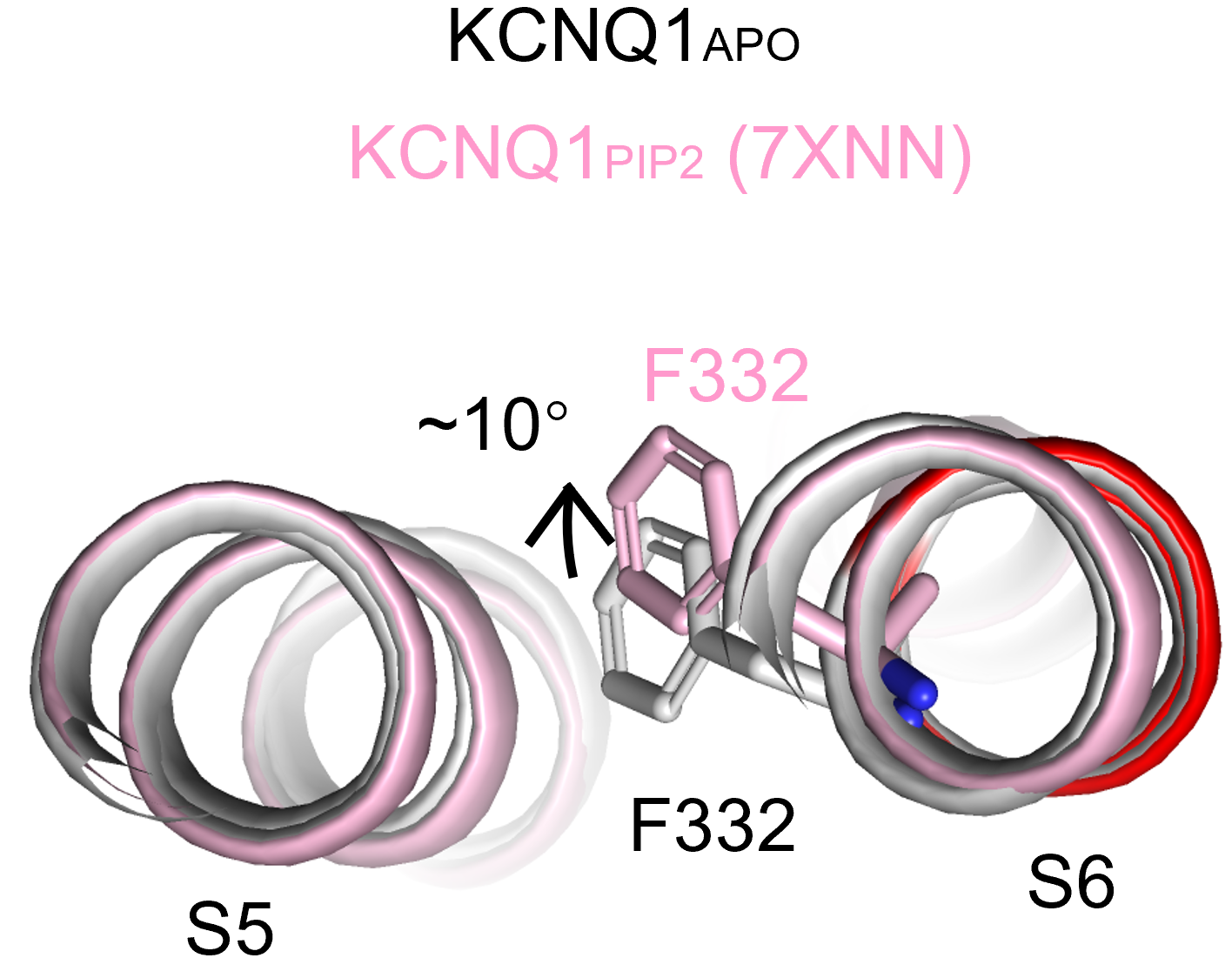

**Figure S14. F332 motion during KCNQ1 channel gating.** From structural comparison between KCNQ1_APO_ and KCNQ1_PIP2_ (PDB: 7XNN ^3^), the F332 residue points to the S5, and undergoes a ~10° clockwise rotation during channel opening.

**Table S1. Cryo-EM data collection, refinement and validation statistics.**

| Structure  EMDB accession code  PDB accession code | KCNQ1-CaMEMDB-64213  9UJ4 | KCNQ1-KCNE1-CaM  EMDB-63935  9U7F | KCNQ1-KCNE1-CaM-PIP2  EMDB-64038  9UC8 |
| --- | --- | --- | --- |
| **Data collection and processing** |  |  |  |
| Magnification | 130,000 | 130,000 | 130,000 |
| Voltage (kV) | 300 | 300 | 300 |
| Electron exposure (e–/Å^2^) | 49.43 | 51.45 | 50.73 |
| Defocus range (μm) | -1.0 ~ -2.0 | -1.0 ~ -2.0 | -1.0 ~ -2.0 |
| Pixel size (Å) | 0.891 | 0.96 | 0.96 |
| Symmetry imposed | *C4* | *C4* | *C4* |
| Initial particle images (#) | 2,443,785 | 1,254,481 | 5,044,889 |
| Final particle images (#) | 245,445 | 152,699 | 168,370 |
| Map resolution (Å)  FSC threshold | 2.59  0.143 | 2.90  0.143 | 3.36  0.143 |
| **Refinement** |  |  |  |
| Initial model used (PDB code) | 6UZZ | 6V00 | 6V01 |
| Model resolution (Å)  FSC threshold | 3.10  0.143 | 3.10  0.143 | 3.90  0.143 |
| **Model composition**  Non-hydrogen atoms  Protein residues  Ligands | 15968  1972  16 | 17224  2128  16 | 15595  2052  11 |
| ***B* factors (**Å**^2^)**  Protein  Ligand | 54.41  35.92 | 60.35  101.46 | 38.08  76.76 |
| r.m.s. deviations  Bond lengths (Å)  Bond angles (°) | 0.004  0.506 | 0.007  1.109 | 0.008  1.175 |
| **Validation**  MolProbity score  Clashscore  Poor rotamers (%) | 1.34  6.22  0.48 | 1.23  2.56  0.00 | 1.99  12.19  0.87 |
| Ramachandran plot  Favored (%)  Allowed (%)  Disallowed (%) | 99.18  0.82  0.00 | 96.85  3.15  0.00 | 94.41  5.39  0.20 |
